# The polyol pathway is a crucial glucose sensor in *Drosophila*

**DOI:** 10.1101/2020.03.16.993170

**Authors:** Hiroko Sano, Akira Nakamura, Mariko Yamane, Hitoshi Niwa, Takashi Nishimura, Masayasu Kojima

## Abstract

A major nutrient source for animals is glucose, which induces transcriptional responses that shift metabolism. Such metabolic adaptation should be appropriately scaled to the ingested glucose levels, and decoupling causes human metabolic diseases. However, the identity of the crucial sensor metabolite(s) that transmit circulating glucose levels to the transcriptional machinery remains elusive. Here we show that the polyol pathway, which converts glucose to fructose via sorbitol, is required for activation of the master metabolic regulator Mondo, the *Drosophila* homologue of MondoA/ChREBP. We demonstrate that under normal nutritional conditions polyol pathway metabolites promote Mondo’s nuclear localization and cause global changes in metabolic gene expression. Polyol pathway mutants block nuclear localization of Mondo and Mondo-mediated gene expression despite intact glycolytic and pentose phosphate pathways. Our results uncover the normal physiological function of this pathway and cast a new light on the adverse effects of high fructose diets in human health.

## Introduction

The accurate sensing of ingested nutrients is vital for organismal survival. Animals optimize metabolism to match the nutrients they detect as being nutritionally available, allowing them to flourish on unstable food supplies. Glucose is the most commonly used carbon source for animals and provides a good example of how animals developed mechanisms that achieve nutritional adaptation. However, glucose is incorporated into a variety of metabolic pathways in cells, which has hampered the identification of the sensor metabolite(s) that inform cells of the availability of glucose.

We previously identified a sugar-responsive peptide hormone, CCHamide-2 (CCHa2) in *Drosophila*. CCHa2 is synthesized mainly in the fat body and the gut in response to glucose ingestion. CCHa2 stimulates brain insulin-producing cells (IPCs) to promote the production of *Drosophila* insulin-like peptides (Dilps) and regulate larval growth (Sano et al., 2015). Despite its important role in coupling systemic growth to glucose availability, the mechanism by which glucose triggers CCHa2 production is unclear. Although acute feeding of abnormally high amounts of sugar (e.g. 20-50% sucrose) increased glucose levels in the hemolymph (Havula et al., 2013; Pasco and Leopold, 2012; Ugrankar et al., 2015), ingestion of glucose at a concentration relevant for what flies consume in nature (~5%) does not change circulating glucose levels significantly, presumably due to high basal glucose concentration in the hemolymph (Mishra et al., 2013; Miyamoto et al., 2012). Thus, glucose does not appear to be sensed directly by cells. These observations prompted us to investigate how glucose uptake is detected in *Drosophila*.

In mammals, MondoA and its paralogue, Carbohydrate responsive element binding protein (ChREBP), are activated in the presence of glucose. Glucose ingestion ultimately promotes MondoA/ChREBP’s nuclear localization and ability to transactivate. A low-glucose inhibitory domain located at the N-terminus of MondoA/ChREBP represses transactivation activity of the protein probably through intramolecular interaction, and this inhibition is released under high glucose conditions by unknown mechanisms (Davies et al., 2010; Li et al., 2006). MondoA/ChREBP are also subject to post-translational modifications. It has been proposed that phosphorylation/dephosphorylation regulate MondoA/ChREBP’s nuclear translocation and DNA-binding (Kabashima et al., 2003; Kawaguchi et al., 2001; Sakiyama et al., 2008); however, phosphomimetic mutants do not recapitulate the glucose-dependent regulation of MondoA/ChREBP (Li et al., 2006; Tsatsos et al., 2008). Thus, the mechanisms by which MondoA/ChREBP is activated by glucose remain a matter of debate. MondoA and ChREBP are preferentially expressed in muscles, and the liver and adipose tissues, respectively. In high glucose conditions, MondoA and ChREBP act together with their obligate partner, Max-like protein X (Mlx), to activate the expression of lipogenic genes thereby storing excess nutrients in the form of lipids. In *Drosophila,* the homologue of MondoA/ChREBP is encoded by a single gene, *Mondo*. Transcriptome analysis has shown that the Mondo-Mlx (also called Bigmax in *Drosophila*) complex induces global changes in metabolic gene expression according to sugar availability (Havula et al., 2013; Mattila et al., 2015). Thus, the downstream effects of Mondo/ChREBP have been well-documented; however, the connection between glucose ingestion and Mondo/ChREBP activation remains elusive.

Sugar metabolites that mediate the transcriptional effects of glucose have been investigated extensively using mammalian cell culture systems. So far, several candidates for MondoA/ChREBP-activating sugars have been reported, including glucose-6-phosphate, xylulose-5-phosphate, and fructose-2,6-bisphosphate (Arden et al., 2012; Dentin et al., 2012; Diaz-Moralli et al., 2012; Iizuka et al., 2013; Kabashima et al., 2003; Li et al., 2010; Peterson et al., 2010; Petrie et al., 2013; Stoltzman et al., 2008). These sugars are synthesized through either glycolysis or the pentose phosphate pathway (PPP), which branches off from the first glycolytic metabolite, glucose-6-phosphate. Thus, cells were thought to detect blood glucose levels by assessing the activities of these two pathways. However, this scenario is undermined by observations that the levels of metabolites in these pathways remain mostly constant after glucose uptake under normal nutritional conditions in which storage sugars accumulate in the cell (Peeters et al., 2017). Storage sugars such as glycogen and trehalose are known to provide buffer action to prevent drastic changes in glucose metabolism; excessive nutrient uptake promotes the conversion of glucose-6-phosphate into glycogen and trehalose, while starvation induces the breakdown of glycogen and trehalose into glucose-6-phosphate and glucose, respectively. Glycolysis is also subject to feedback activation or inhibition by downstream metabolites (Mor et al., 2011). Furthermore, endocrine regulation by insulin and glucagon could disrupt a linear response of glycolysis to glucose uptake into the cell. These findings support the hypothesis that unknown pathways are involved in the detection of blood glucose levels.

In this study, we show that *Drosophila* Mondo is required for glucose-induced *CCHa2* expression. We further show that activation of Mondo upon glucose ingestion is regulated by the polyol pathway, an evolutionary conserved metabolic pathway in which glucose is reduced to sorbitol by aldose reductase, then converted to fructose by sorbitol dehydrogenase (Hers, 1956). Polyol pathway metabolites promote the nuclear localization of Mondo in the *Drosophila* fat body, an organ analogous to mammalian liver and adipose tissues. Furthermore, transcriptional regulation of most Mondo/Mlx-target genes, including metabolic genes, was affected in mutants lacking polyol pathway enzymes. Hyper activation of the polyol pathway has been implicated in the exacerbation of disease, such as diabetic complications in humans and cancer metastasis in cell culture (Brownlee, 2001; Lorenzi, 2007; Schwab et al., 2018). However, an understanding of the physiological roles of the polyol pathway has remained largely elusive, hindering a consideration of how the polyol pathway exerts pathogenic effects. Our study clarifies the long-standing question about the normal physiological function of the polyol pathway and provides a basis for examining the mechanisms of polyol pathway-associated disease aggravation.

## Results

### Sugar-dependent *CCHa2* expression is mediated by Mondo

To dissect the mechanisms of glucose detection, we used *CCHa2* expression as a test case. Transcriptome analysis using whole larvae had identified *CCHa2* as a Mondo-Mlx target gene (Mattila et al., 2015). Since *CCHa2* is preferentially expressed in the fat body (Chintapalli, 2010; Sano et al., 2015), we examined whether Mondo regulates *CCHa2* tissue autonomously. Larvae were raised in regular fly food containing 10% glucose and *CCHa2* mRNA levels were measured using quantitative RT-PCR (RT-qPCR). As shown in **Figure 1A**, fat body-specific knockdown of Mondo significantly decreased *CCHa2* expression (**Figure 1A**), indicating that Mondo functions autonomously in this organ for normal *CCHa2* expression. Fat body-specific knockdown of *sugarbabe* (*sug*), a transcription factor acting downstream of Mondo (Mattila et al., 2015), also reduced *CCHa2* mRNA levels. These results indicate that Mondo controls *CCHa2* expression either directly or indirectly through *sug* in fat body cells.

**Figure 1.**
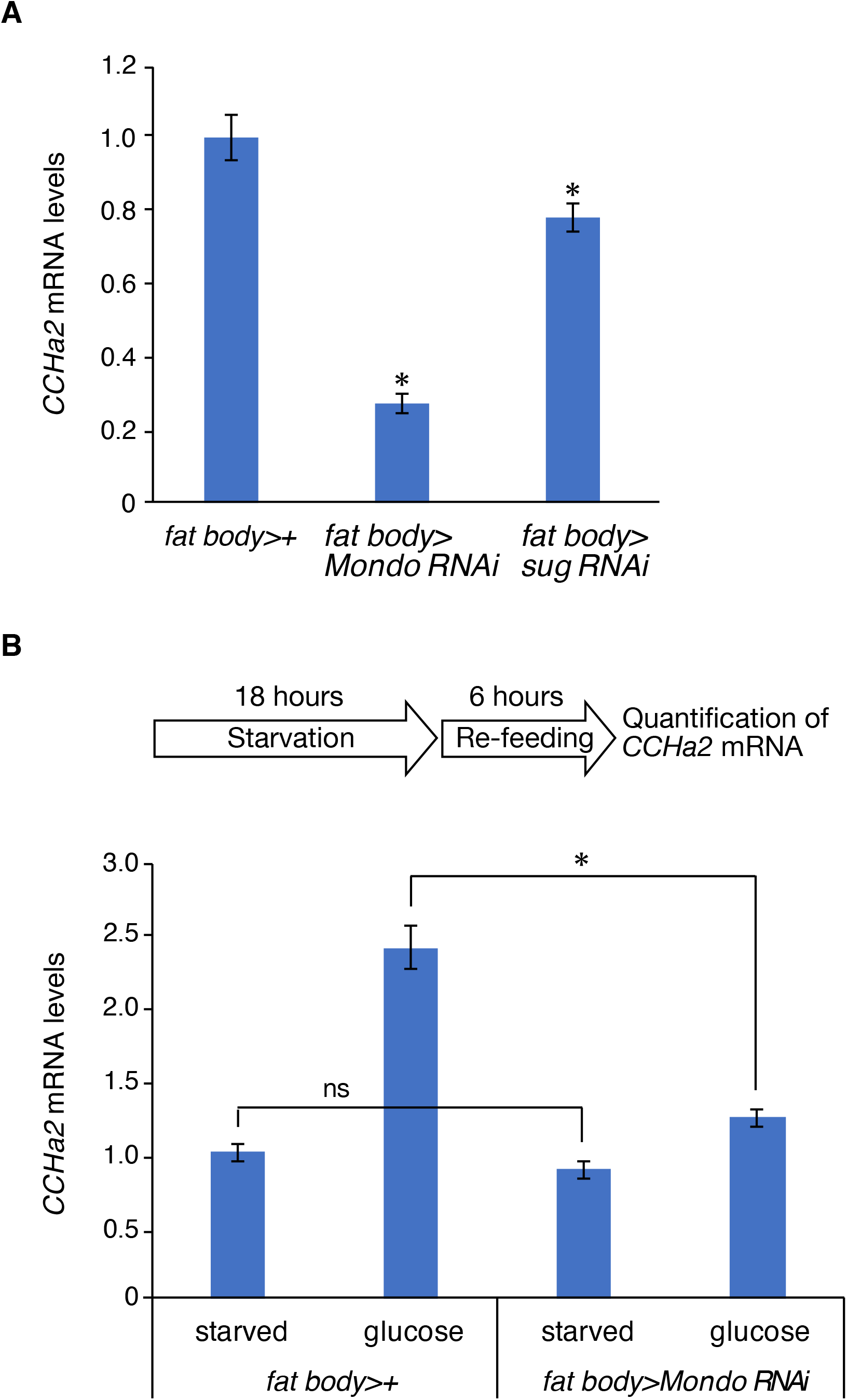
Mondo mediates sugar-dependent *CCHa2* expression. (A) *CCHa2* mRNA levels in third-instar larvae (72 hours AEL) were measured by RT-qPCR. Fat body-specific knockdown of *Mondo* and *sug* was conducted using the *cg-GAL4* driver in combination with *UAS-Mondo RNAi* and *UAS-sug RNAi*, respectively. Fat body-specific knockdown of *Mondo* and *sug* significantly reduced *CCHa2* mRNA levels. 10 larvae per batch, n = 3 batches for all genotypes. (B) Induction of *CCHa2* expression by glucose was tested in third-instar larvae. Larvae starved for 18 hours were re-fed with 10% glucose for 6 hours, and *CCHa2* mRNA levels in whole animals were measured by RT-qPCR. Fat body-specific knockdown of *Mondo* significantly reduced glucose-dependent *CCHa2* expression. 10 larvae per batch, n = 3 batches for all genotypes. Histograms show mean ± SE. *p < 0.05, ns, not significant.

We then examined the role of Mondo in the induction of *CCHa2* expression under fluctuating nutritional conditions. In this assay, early third-instar larvae were starved for 18 hours on water agar plates. The larvae were then re-fed with glucose for 6 hours, and *CCHa2* mRNA levels were quantified. *CCHa2* expression was elevated by glucose ingestion in control larvae, while knockdown of Mondo in the fat body significantly reduced glucose-induced *CCHa2* expression (**Figure 1B**). These results show that Mondo is required for sugar-dependent induction of *CCHa2* expression in the fat body.

### The polyol pathway is required for Mondo-mediated *CCHa2* expression

We next sought to reveal how glucose ingestion led to Mondo-mediated *CCHa2* expression. Mammalian cell culture studies have identified glucose-6-phosphate, xylulose-5-phophate, and fructose-2, 6-biphosphate as Mondo-activating metabolites, which can be produced through either glycolysis or PPP (Arden et al., 2012; Dentin et al., 2012; Diaz-Moralli et al., 2012; Iizuka et al., 2013; Kabashima et al., 2003; Li et al., 2010; Peterson et al., 2010; Petrie et al., 2013; Stoltzman et al., 2008). In our previous studies, we demonstrated that re-feeding of starved larvae with glucose, sorbitol, or fructose was able to induce *CCHa2* expression (**Figure 2A**) (Sano, 2015; Sano et al., 2015). Glucose and fructose are metabolized by glycolysis and PPP; however, sorbitol is not metabolized by either pathway, suggesting that other metabolic pathway(s) are involved in *CCHa2* expression. The Kyoto Encyclopedia of Genes and Genomics (KEGG) database indicates that the polyol pathway, in which glucose is reduced to sorbitol by aldose reductase (EC: 1.1.1.21) then oxidized to fructose by sorbitol dehydrogenase (EC: 1.1.1.14), is the only pathway that can metabolize sorbitol in *Drosophila* (**Figure 2B**) (Kanehisa and Goto, 2000; Kanehisa et al., 2019). This suggests that the metabolites sensed by cells to induce *CCHa2* could be a product of the polyol pathway.

**Figure 2.**
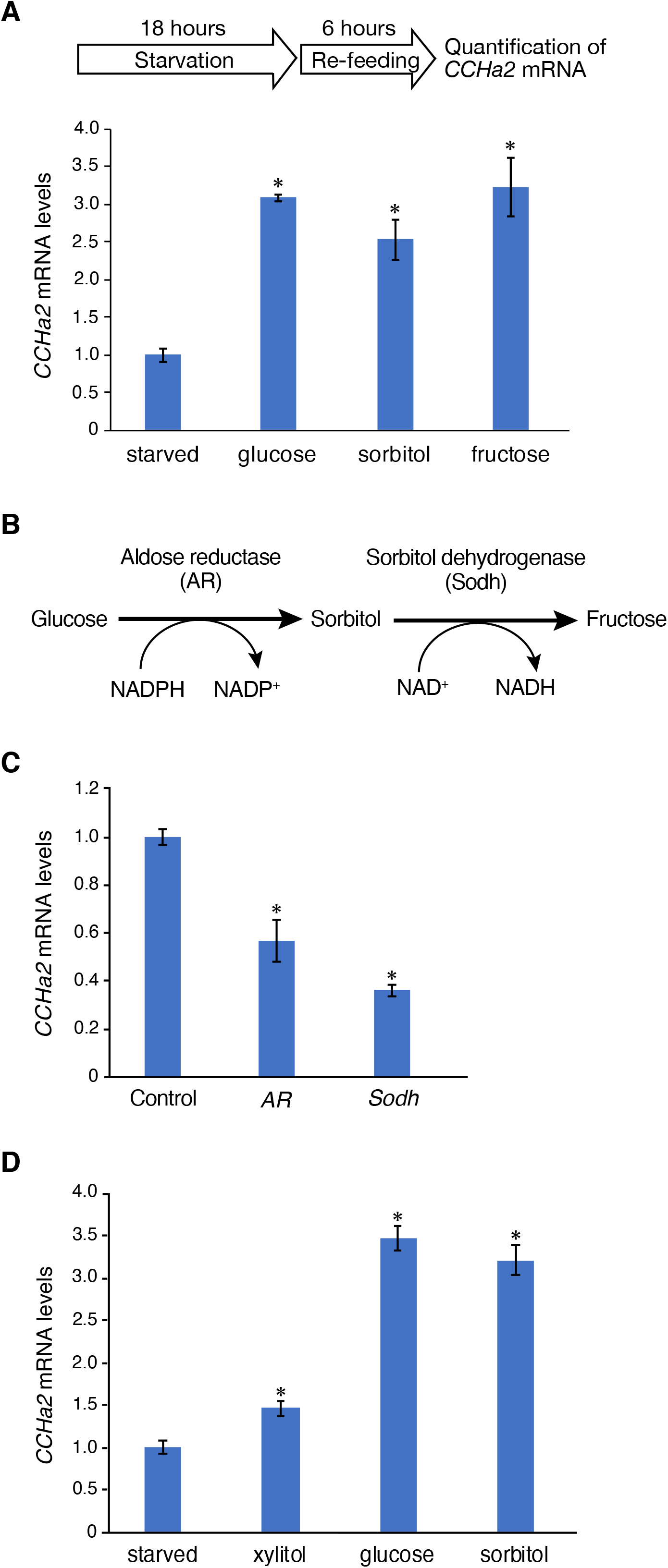
The polyol pathway is required for Mondo-mediated *CCHa2* expression. (A) Effects of different sugars on *CCHa2* expression in wild-type larvae. Third-instar larvae (72 hours AEL) starved for 18 hours were re-fed for 6 hours with a 10% solution of the indicated sugars, and *CCHa2* mRNA levels in whole animals were measured by RT-qPCR. (B). The polyol pathway consists of two steps catalyzed by aldose reductase then sorbitol dehydrogenase. The first step generates sorbitol from glucose, the second fructose from sorbitol. (C) *CCHa2* mRNA levels in the third-instar larvae were measured by RT-qPCR. A significant reduction in *CCHa2* expression was observed in *AR* and *Sodh* mutant larvae raised on a regular diet containing 10% glucose. (D) The effects of xylitol on *CCHa2* expression were compared to those of glucose and sorbitol in the third-instar larvae (72 hours AEL). Larvae starved for 18 hours were re-fed with 10% xylitol, glucose, or sorbitol for 6 hours, and *CCHa2* mRNA levels in whole animals were measured by RT-qPCR. 10 larvae per batch, n = 3 batches for all genotypes. Histograms show mean ± SE. *p < 0.05.

To test this hypothesis, we genetically blocked the activity of the polyol pathway and examined whether *CCHa2* induction was affected. The KEGG database predicts seven aldose reductase (AR) genes in the *Drosophila* genome. Among those, we targeted both *CG6084* and *CG10638,* creating double mutants. There are two sorbitol dehydrogenase genes, *Sodh-1* and *Sodh-2*. Strong sequence similarities between them suggest redundant functions, thus we mutated both genes. To create each of these double mutant lines we designed single guide RNAs (sgRNAs) targeting the coding regions and introduced small deletions using the CRISPR/Cas9 system (**Figures S1–S3**). This caused a frame shift and a premature termination in the translated proteins, blocking production of the catalytic portion of these enzymes. To validate this conclusion we conducted gas chromatography mass spectrometry (GC/MS) analysis of the larval hemolymph. We detected a reduction of sorbitol in the *CG6084*, *CG10638* double mutants, hereafter named *AR* mutants (**Figure S4A**). We also observed an accumulation of sorbitol in the *Sodh-1*, *Sodh-2* double mutants, hereafter named *Sodh* mutants (**Figure S4A**). We first used these mutants to examine *CCHa2* expression in larvae under normal culture conditions. Mutant larvae were raised on regular fly food containing 10% glucose and *CCHa2* mRNA levels in the third-instar larvae (72 hours AEL) were quantified. The results show that *CCHa2* mRNA levels were significantly reduced in the *AR* mutants (**Figure 2C**). A similar reduction in *CCHa2* expression was observed in the *Sodh* mutants (**Figure 2C**). Notably, *CCHa2* mRNA levels in these mutants were reduced even though genes involved in glycolysis and PPP were intact. These results indicate that the polyol pathway plays a crucial role in *CCHa2* expression under normal feeding conditions.

In addition to catalyzing the conversion of glucose to fructose, *AR* and *Sodh* are predicted to transform xylose to xylulose via xylitol. AR converts xylose to xylitol, which is then oxidized by Sodh to produce xylulose (**Figure S4B**; KEGG pathway database). GC/MS analysis of the hemolymph from *AR* and *Sodh* mutant larvae showed metabolic phenotypes as predicted; xylitol was reduced in *AR* mutants whereas it was increased in *Sodh* mutants (**Figure S4B**). This raised the possibility that xylulose and its metabolites could also be involved in *CCHa2* expression. Therefore, we tested the effects of xylitol ingestion on *CCHa2* expression and found that the levels of *CCHa2* expression induced by re-feeding starved larvae with xylitol was much lower than those seen upon glucose and sorbitol re-feeding (**Fig. 2D**). Therefore, the contribution of this pathway to regulate *CCHa2* expression would be minor.

### The polyol pathway is required for nuclear localization of Mondo in the fat body

It has been shown that the activity of mammalian MondoA and ChREBP is regulated by its nuclear localization and transactivation. In order to examine how the polyol pathway regulates the activity of *Drosophila* Mondo, we knocked-in the Venus fluorescent protein at the C-terminal end of the Mondo coding region using the CRISPR/Cas9 system (**Figure S5**). The *Mondo::Venus* knockin flies were homozygous viable and fertile, indicating that the *Venus* insertion does not interfere with Mondo function. We observed an intracellular localization of the Mondo::Venus fusion protein in *ex vivo* culture of fat bodies dissected from third-instar larvae raised on normal food. In Schneider’s medium that contains 5 mM glucose, 5.4% of Mondo::Venus was localized in the nuclei of the wild-type fat body cells (**Figures 3A and 3B**). Upon the addition of 55 mM glucose, sorbitol, or fructose into the medium, the percentage of nuclear Mondo::Venus increased to 15.6%, 13.0%, or 16.4%, respectively (**Figures 3A and 3B**). In contrast, in the *AR* mutants, localization of Mondo::Venus was unchanged upon the addition of glucose, while nuclear Mondo::Venus was increased by sorbitol or fructose administration (**Figures 3A and 3C**). In the *Sodh* mutants, only fructose was able to promote the nuclear localization of Mondo::Venus (**Figures 3A and 3D**). Thus, the addition of fructose or sugars that can be converted to fructose through the polyol pathway is sufficient to promote the nuclear localization of Mondo.

**Figure 3.**
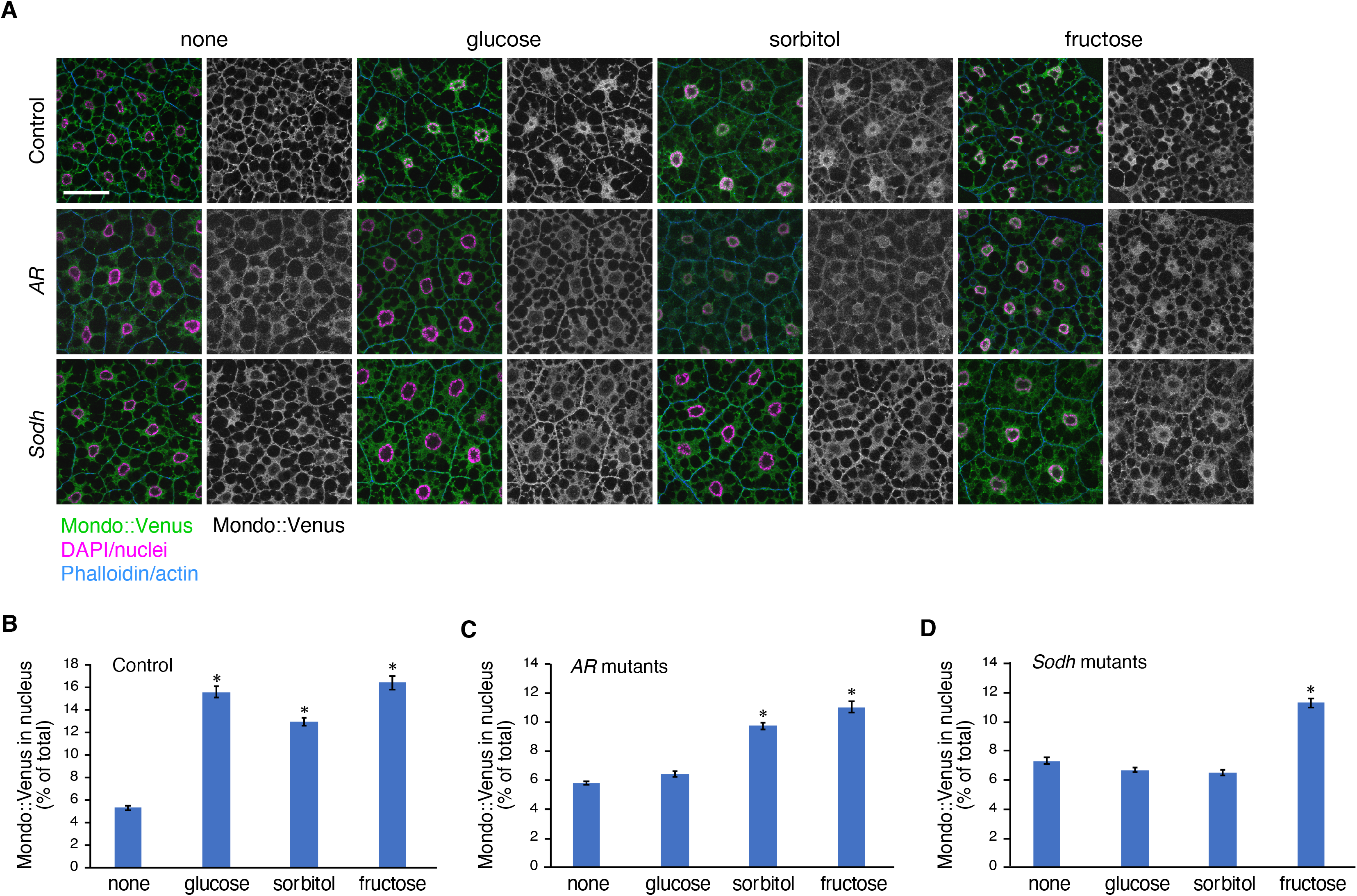
The polyol pathway is required for nuclear localization of Mondo in the fat body. (A) Fat bodies from the Mondo::Venus knockin line were cultured *ex vivo* in Schneider’s medium supplemented with 55 mM glucose, sorbitol, or fructose for 15 minutes. After culture, the fat bodies were fixed and stained with the following markers: anti-GFP antibody for Mondo::Venus (green), DAPI for nuclei (magenta), and Rhodamine-conjugated phalloidin for cortical actin at the cell membrane (blue). Multi-channel images and single channel images showing Mondo::Venus signals are shown. (B-D) The percentage of nuclear Mondo::Venus out of the total Mondo::Venus in a cell was quantified on the images. The quantification reveals that the amount of nuclear Mondo::Venus was significantly increased upon the addition of glucose, sorbitol, and fructose in wild type (A, B). Nuclear localization of Mondo::Venus was promoted by the addition of sorbitol and fructose but not glucose in *AR* mutants (A, C). Nuclear localization of Mondo::Venus was only promoted by fructose in *Sodh* mutants (A, D). Images of 44 to 56 fat body cells per experiment were used for quantification. Scale bar represents 50 μm. Histograms show mean ± SE. *p < 0.05.

### The polyol pathway regulates the majority of Mondo/Mlx-target genes

Since Mondo and its partner Mlx are pivotal in the regulation of metabolic genes (Mattila et al., 2015), we wished to test whether the polyol pathway allows coupling of sugar ingestion to global transcriptional alteration through Mondo. We therefore explored the role of the polyol pathway on the expression of Mondo target genes using RNA-seq. We starved wild-type third-instar larvae (72 AEL) for 18 hours, and then re-fed them with either glucose or sorbitol for 6 hours; we examined RNA seq libraries created from these larvae for the levels of sugar-responsive Mondo/Mlx*-*target genes observed in the datasets reported by Mattila *et al.* (2015) (Mattila et al., 2015). We designed this experiment to distinguish the effects of polyol pathway metabolites from those of glycolysis and PPP which are also upstream of Mondo (Richards et al., 2017). Given that sorbitol is metabolized only through the polyol pathway, polyol pathway metabolites would be selectively increased in sorbitol-fed larvae, whereas metabolites of all three pathways would be increased in glucose-fed larvae. Thus, the transcriptome of sorbitol-fed larvae is expected to reveal genes responsive to the polyol pathway, and that of glucose-fed larvae represents genes responsive to glycolysis, PPP and the polyol pathway.

In order to clarify the contribution of the polyol pathway in Mondo-mediated gene expression, we compared transcriptional changes observed upon sorbitol feeding with those seen upon glucose feeding. As shown in **Figure 4A**, the RNA-seq data shows good agreement between glucose-dependent and sorbitol-dependent transcriptional changes [correlation coefficient (*r*) = 0.92]. We identified 504 glucose-responsive Mondo/Mlx-target genes. Of those, 388 genes (77.0%) were also responsive to sorbitol (**Figure 4B**). These data demonstrate that the polyol pathway can solely regulate Mondo/Mlx-target genes. Transcriptome changes upon sorbitol feeding were almost abolished in the *Sodh* mutants (*r* = 0.16), confirming the essential function of the polyol pathway in sorbitol-dependent transcriptional regulation (**Figure 4C**). In order to confirm the involvement of the polyol pathway in Mondo/Mlx-mediated gene expression, we investigated if the loss of the regulation of Mondo/Mlx-target genes in the *Sodh* mutants is restored by feeding the larvae with fructose. Although glucose-regulated Mondo/Mlx-target genes displayed little change in their expression upon feeding of the *Sodh* mutant larvae with sorbitol (*r* = 0.19, **Figure 4D**), the expression pattern in mutant larvae was mostly restored by fructose ingestion (*r* = 0.54, **Figure 4E**). To check if differences in sugar-dependent gene expression between wild type and *Sodh* mutants could be derived from variations in their genetic background, we examined the transcriptomes of wild-type and *Sodh* mutant larvae under starved conditions. They correlated well (*r* = 0.94, **Figure 4F**), arguing that the observed differences upon sugar feeding are due to the induced mutation in the polyol pathway. These results reveal a substantial contribution of the polyol pathway to Mondo/Mlx-dependent transcriptional regulation.

**Figure 4.**
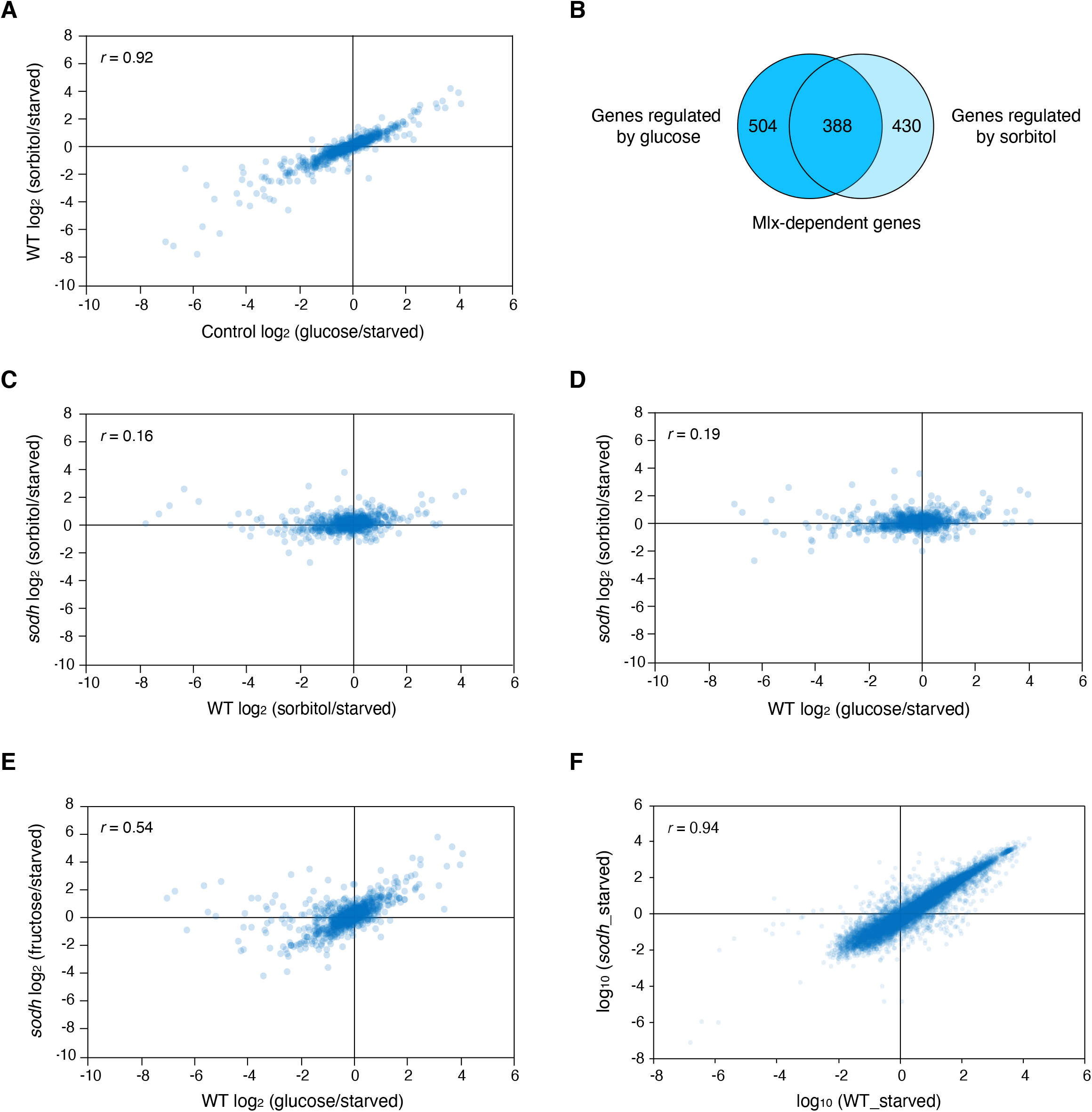
The polyol pathway regulates the majority of Mondo/Mlx-target genes. Transcriptome analysis of sugar-fed larvae. Wild-type or *Sodh* mutant third-instar larvae (72 hours AEL) starved for 18 hours were re-fed for 6 hours with 10% solution of indicated sugars. Transcriptomes of whole animals were obtained using RNA-seq and the expression patterns of Mlx-target genes are plotted in the graph. (A, B) A comparison of the Mlx-target gene expression between the glucose-fed and sorbitol-fed larvae. Good correlation (*r* = 0.92) was observed. 504 Mlx-target genes were regulated by glucose. Of those, 430 genes (77.0%) were regulated by sorbitol. (C) A comparison of the Mlx-target gene expression between sorbitol-fed wild-type and sorbitol-fed *Sodh* mutant larvae. The changes in gene expression observed in wild-type were abolished in *Sodh* mutants, indicating that gene expression caused by sorbitol depends on the activity of the polyol pathway (r=0.16). (D. E) The loss of sorbitol-induced Mlx-target gene expression in *Sodh* mutants was partially rescued by fructose (r=0.19, 0.54 respectively). (F) Transcriptomes of starved wild-type and *Sodh* mutant larvae were identified using RNA-seq. The transcriptomes of these animals were well correlated (*r* = 0.94), indicating that there is no significant difference in gene expression between wild type and *Sodh* mutants under starved conditions. 30 larvae per batch, n = 3 batches for all genotypes.

It has been reported that Mondo and Mlx regulate genes involved in glycolysis/gluconeogenesis, PPP, fatty acid biosynthesis, and glutamate and serine metabolism (Mattila et al., 2015). We observed similar expression patterns of these metabolic genes between glucose-fed and sorbitol-fed wild-type larvae (**Figure 5**), indicating that the polyol pathway can solely regulate most of the Mondo/Mlx-target metabolic genes. In sorbitol-fed *Sodh* mutant larvae in which glycolysis, PPP, and the polyol pathway are expected to remain at levels akin to starvation conditions, the pattern was altered. In contrast, the pattern was largely restored in fructose-fed *Sodh* mutant larvae in which the downstream output of the polyol pathway is restored. In particular, the Mondo/Mlx-target genes identified in (Mattila et al., 2015) (indicated in red and green in **Figure 5**) were remarkably rescued in fructose-fed *Sodh* mutant larvae. These results demonstrate that the polyol pathway is the principal metabolic pathway involved in glucose-dependent activation of Mondo, required to induce global changes in metabolic gene expression.

**Figure 5.**
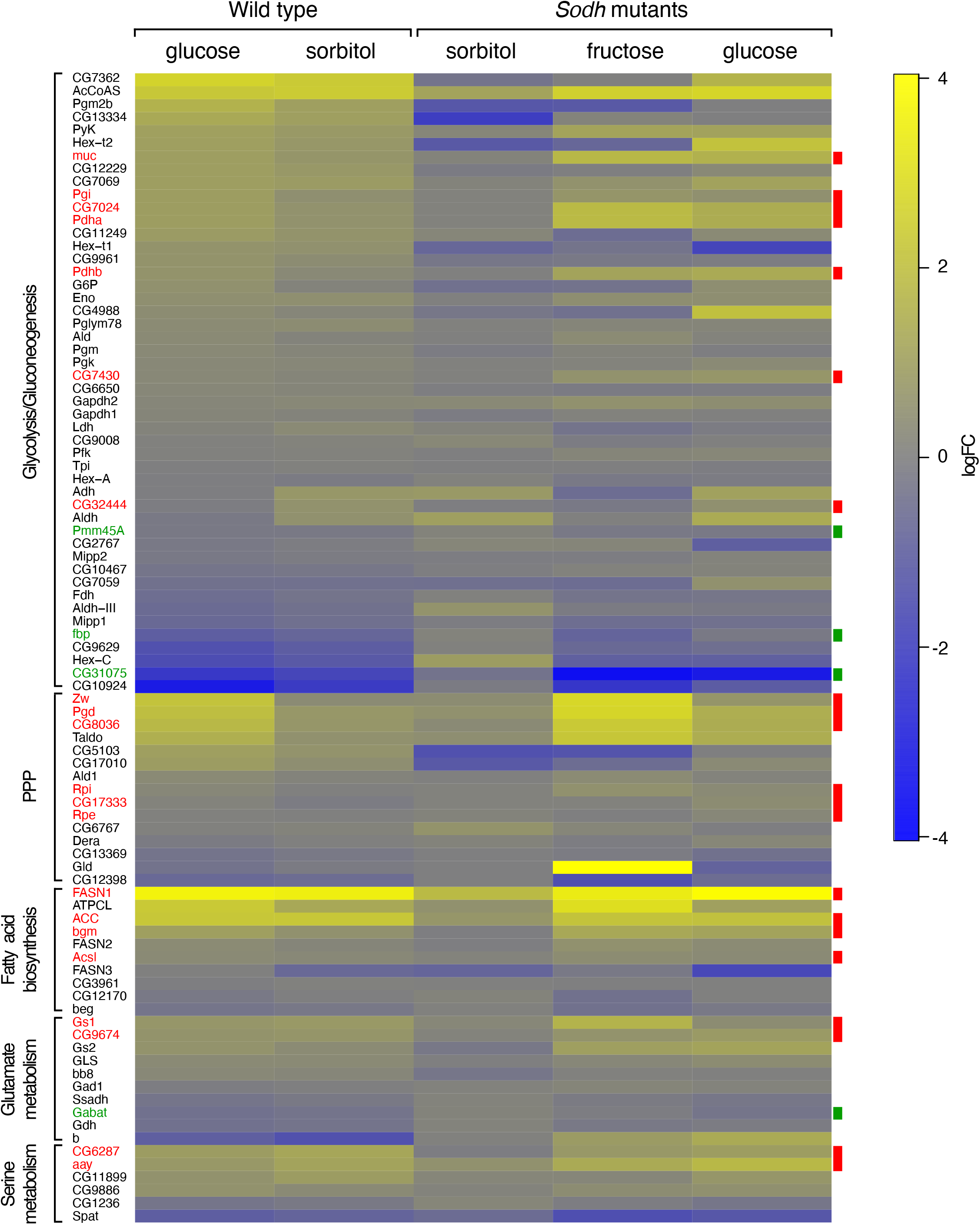
The polyol pathway regulates key metabolic enzymes acting downstream of Mondo/Mlx. The expression pattern of genes involved in glycolysis/gluoneogenesis, PPP, fatty acid biosynthesis, glutamate and serine metabolism in the starvation/re-feeding assay. The names of genes upregulated by Mondo/Mlx are indicated in red, and those downregulated by Mondo/Mlx are indicated in green. The results of five experiments (genotype/fed sugar) are shown. Changes in gene expression are represented as colors: yellow to indicate upregulation, and blue to indicate downregulation. Feeding glucose and sorbitol produced similar changes in gene expression. The pattern was altered in sorbitol-fed *Sodh* mutants. The expression pattern was partially restored by feeding mutant larvae with fructose or glucose. Mondo/Mlx-target genes showed a remarkable rescue.

### Mechanisms of glucose detection in different nutritional conditions

The above results show that the polyol pathway is important for Mondo-mediated transcription: in normally-fed larvae even in the presence of normal glycolysis and PPP activities, the polyol mutations showed a significant reduction in *CCHa2* expression (**Figure 2C**) and glucose-dependent nuclear localization of Mondo (**Figure 3**). However, we obtained data that indicated the presence of a more complicated regulatory landscape. When using starved *Sodh* mutant larvae, the ingestion of glucose still induced *CCHa2* expression normally even in the absence of the polyol pathway (**Figure 6A**). These larvae even display regulation of a considerable number of Mondo/Mlx-target genes, including metabolic genes (**Figures 5 and 6B**). Intriguingly, the 18-hour-starvation we used prior to the re-feeding causes complete consumption of glycogen in the fat body (Matsuda et al., 2015). The depletion of storage sugar in animal cells is known to lead to the loss of homeostasis of glycolysis and PPP. Therefore, distinct glucose-sensing systems seem to operate under different nutritional conditions. Taken together, we propose that the polyol pathway is the principal glucose-sensing pathway under normal nutritional conditions, and that glycolysis and PPP can contribute to glucose sensing in starved animals that lack metabolic homeostasis.

**Figure 6.**
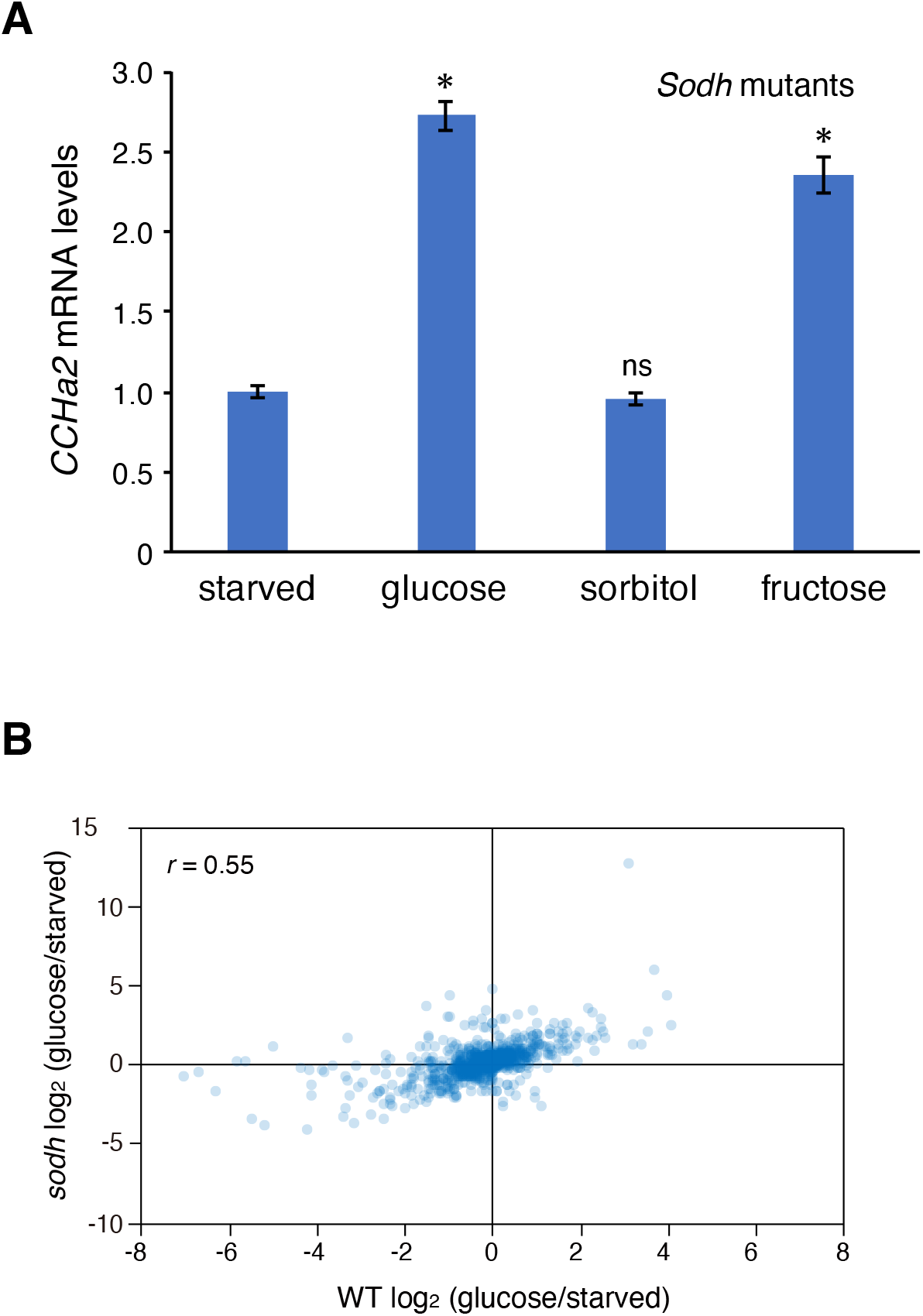
The polyol pathway is not essential for glucose detection under starved conditions. (A) Effects of different sugars on *CCHa2* expression in *Sodh* mutant larvae. Third-instar larvae starved for 18 hours were re-fed for 6 hours with 10% solution of indicated sugars, and *CCHa2* mRNA levels in whole animals were measured by RT-qPCR. Glucose and fructose but not sorbitol significantly promoted *CCHa2* expression. (B) A comparison of the Mlx-target gene expression between glucose-fed wild-type and glucose-fed *Sodh* mutant larvae. Considerable numbers of glucose-regulated genes detected in wild-type were regulated by glucose in *Sodh* mutants (*r* = 0.55). 10 larvae per batch, n = 3 batches for all genotypes. Histograms show mean ± SE. *p < 0.05, ns, not significant, in (A). 30 larvae per batch, n = 3 batches in (B).

## Discussion

Direct sugar detection by cells is thought to be a sugar sensing module present in lower animals without endocrine systems that is conserved in higher organisms with multiple layers of metabolic homeostasis, including humans. Such sugar perception regulates various physiological responses, particularly metabolism, allowing organisms to adapt to nutritional changes primarily through transcriptional regulation. The existence of sensor metabolite(s) that activate the sugar-responsive transcription factors has long been anticipated. Several candidate metabolites have been identified in different experimental conditions, yet the true active agent has remained a matter of debate. Here, we show that the polyol pathway is crucial for the activation of Mondo, a master transcription factor that regulates various metabolic genes. Our results demonstrate that the polyol pathway is the principal glucose sensing pathway under normal nutritional conditions. We additionally provide evidence that glycolysis and PPP also contribute to glucose sensing particularly under starved conditions. Although the polyol pathway has been implicated in the aggravation of diabetes and cancer metastasis, the underlying mechanisms are poorly understood. Our study provides a holistic view of glucose sensing by animal cells and a new perspective on polyol pathway-related disease mechanisms.

### The significance of the polyol pathway in sugar sensing

To enable proper organismal adaptation to ingested sugars, the concentration of the sensor metabolite(s) and the activity of the sensor metabolic pathway(s) need to correlate with the levels of glucose in the body fluid. The polyol pathway appears to fulfill these conditions. First, the polyol pathway is the most upstream glucose-metabolizing pathway. Glucose flows into the polyol pathway before being metabolized to glucose-6-phosphate, which is consumed through glycolysis and PPP (**Figure 7**). Second, the polyol pathway would be less affected by storage sugars. Glycogen, the major carbohydrate storage form in animal cells, is converted reversibly into glucose-6-phosphate according to nutrient status of the cell (**Figure 7**). Trehalose is also synthesized from glucose-6-phosphate under eutrophic conditions and is hydrolyzed into glucose under starved conditions. Both of these adjustments to different nutritional states maintain constant glucose-6-phosphate levels, thereby aiding the metabolic homeostasis of glycolysis and PPP regardless of the availability of glucose (Peeters et al., 2017). Third, no feedback control on the polyol pathway has been reported. This is a sharp contrast to glycolysis, in which several enzymes are subject to feedback control by downstream metabolites. For example, phosphofructokinase 1 (PFK1), a rate-limiting enzyme of glycolysis that phosphorylates fructose-6-phosphate, is controlled by several downstream metabolites such as ATP, citrate, ADP, AMP, cAMP and fructose-2,6-bisphosphate (Mor et al., 2011). Furthermore, phosphofructokinase 2 (PFK2), another kinase for fructose-6-phosphate is regulated by insulin- and glucagon-mediated endocrine signaling (Mor et al., 2011). These observations suggest that the polyol pathway could exhibit a linear response to glucose levels in the body fluid better than glycolysis and PPP under normal feeding conditions. Therefore, it is conceivable that the polyol pathway acts as a major glucose-sensing pathway under conditions in which homeostasis of major glucose metabolic pathways is maintained by storage sugars and feedback control. Our results also show that glycolysis and PPP act as additional glucose-sensing pathways in starved animals (**Figure 6**). These observations suggest that multiple glucose sensors exist and function in different nutritional conditions. Having multiple glucose sensing systems would be beneficial for cells and organisms for their adaptation to distinct and fluctuating nutritional conditions.

**Figure 7.**
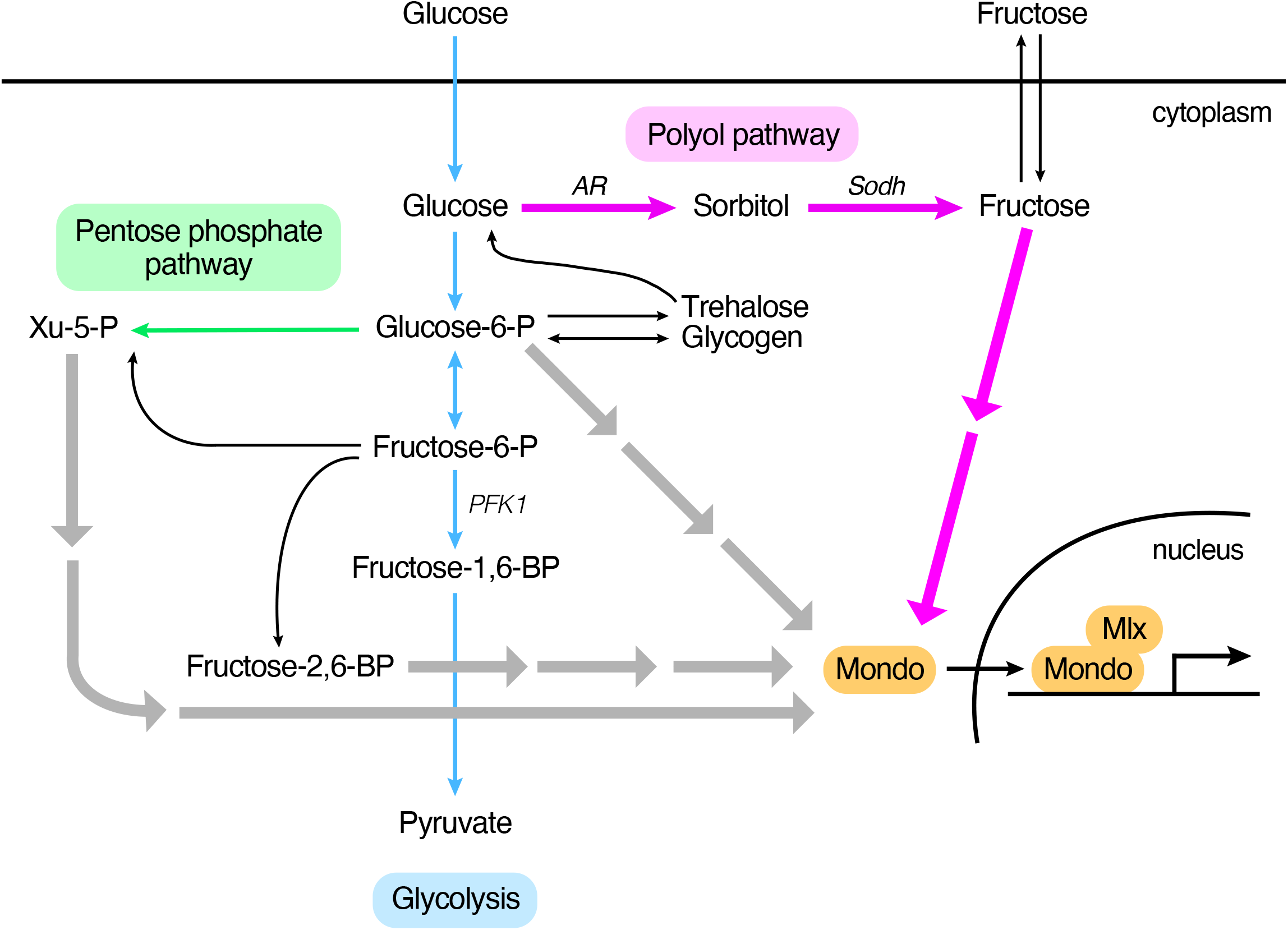
Metabolic pathways leading to Mondo activation. Glucose-6-phosphate, xylulose-5-phosphate, and fructose 2,6-biphosphate were identified previously as MondoA or ChREBP-activating metabolites in mammalian cell culture systems (gray arrows). These metabolites are generated during glycolysis (blue arrows) or PPP (green arrow). Our study revealed that the polyol pathway (magenta arrows) has a significant contribution to Mondo-dependent transcription in *Drosophila* (magenta arrows).

Our results suggest that fructose and fructose derivatives are good candidates for the Mondo-activating sensor metabolite(s) during polyol pathway-mediated glucose sensing. In larval stages, *AR* is expressed ubiquitously, and *Sodh* is preferentially expressed in the fat body and gut (Graveley, 2011). Therefore, the sensor metabolite(s) are likely to be synthesized in these organs and secreted to the hemolymph from which they signal to cells in other organs. It has been shown that the concentration of circulating fructose is acutely elevated after glucose ingestion, probably due to the low basal concentration of fructose in the hemolymph (Miyamoto et al., 2012). Therefore, the conversion of a part of ingested glucose into fructose could be advantageous to allow glucose detection, especially in hyperglycemic animals such as insects. Given that the GS/MS results show that the concentration of fructose in the hemolymph varies even between control lines (*Oregon-R* and *white*) (**Figure S4**), we suggest that cells recognize the dynamics of sensor metabolite(s) rather than their absolute hemolymph concentration. Hyperglycemic conditions are also observed in the mammalian liver to which dietary glucose is carried directly from the small intestine. In contrast, the liver contains only little fructose, as most dietary fructose is cleared in the small intestine by ketohexokinase (Khk) (Jang et al., 2018). In humans, *AR* genes*, AKR1B1* and *AKR1B10*, are weakly expressed in the liver, and the *Sodh gene, SORD,* is strongly and preferentially expressed in the liver (Fagerberg et al., 2014). Thus, the polyol pathway could also function for proper sensing of sugar ingestion in the mammalian liver.

### Tissue specificity of sugar sensing mechanism

It has been shown that adult flies can sense fructose through a narrowly tuned fructose receptor, Gr43a (Miyamoto et al., 2012). Gr43a in the taste sensillae detects external fructose, while Gr43a in the brain senses internal fructose. Since Gr43a is broadly expressed in larval tissues including the fat body (Chintapalli, 2010), it could contribute to sugar sensing in this organ. However, Gr43a is dispensable for sugar-dependent *CCHa2* expression, since *CCHa2* expression occurred normally in *Gr43a* mutant larvae in response to glucose (**Figure S6**). In general, a gustatory receptor is depolarized upon ligand binding, leading to rapid neuronal responses. Indeed, *Gr43a*-mediated fructose preference in larvae is established in minutes (Mishra et al., 2013). The lack of Gr43a-mediated sugar sensing in the fat body suggests that such a rapid response to the nutritional status is unnecessary in the storage organs, and that different nutrient sensing operates in different tissues.

### Interplay between direct and indirect sugar sensing systems

In higher organisms, the response to dietary glucose is a combined effect of direct sugar sensing by cells and the subsequent modification of this cellular sugar sensing by endocrine signaling. An interplay between direct and indirect mechanisms is well documented in mammalian pancreatic β cells. In these cells, glucose metabolism inhibits ATP-sensitive potassium (K_ATP_) channels and opens voltage-dependent calcium channels, which results in the exocytosis of the insulin-containing granules (Straub and Sharp, 2002). Similar mechanisms appear to operate in the *Drosophila* insulin producing cells (IPCs) in the brain, as glucose can induce a Ca^2+^ influx which is also seen upon the pharmaceutical inhibition of K_ATP_ channels (Fridell et al., 2009). However, recent studies have identified peripheral tissues including the fat body, gut, and prothoracic gland as being able to regulate *Drosophila* insulin like peptide (Dilp) secretion (Agrawal et al., 2016; Delanoue et al., 2016; Kim and Neufeld, 2015; Koyama and Mirth, 2016; Rajan and Perrimon, 2012; Sano et al., 2015; Sun et al., 2017), illuminating important roles for non-cell autonomous regulation of IPCs in Dilp secretion. However, the mechanisms for sugar detection in these peripheral tissues remained elusive. We show here that the polyol pathway-Mondo axis allows direct sugar sensing in larval tissues, and that *CCHa2* is one of polyol-Mondo target genes in the fat body. Given that fat body-derived CCHa2 directly stimulates IPCs for the production and secretion of Dilps (Sano et al., 2015), CCHa2 links polyol-Mondo-mediated direct and Dilp-mediated indirect sugar sensing systems, forming a regulatory circuit for glucose homeostasis. Interestingly, another larval sugar-responsive Dilp regulator, Adipokinetic hormone (AKH), was not induced by the polyol-Mondo pathway. AKH secretion is promoted by trehalose (Kim and Neufeld, 2015). Trehalose is a major hemolymph sugar that serves as an energy source in most insects, thus high hemolymph trehalose would be recognized by cells as a chronic high sugar condition. The lack of polyol-Mondo-mediated regulation of AKH expression suggests that different mechanisms are used to detect acute and chronic sugar uptake for proper maintenance of sugar homeostasis.

### Insights into fructose-induced pathogenic mechanisms

Humans find fructose sweeter than glucose at equal concentrations. High-fructose corn syrup has been used in artificially sweetened foods since the 1970s, and fructose consumption has increased drastically over the past decades. Epidemiological studies have shown that the increase in fructose consumption correlates with that in metabolic diseases including obesity, fatty liver, and nonalcoholic fatty liver disease (Bray et al., 2004; Lim et al., 2010; Marriott et al., 2009). Experimental studies have revealed that fructose administration to the cell elevates lipid accumulation better than glucose does (Stanhope et al., 2009; Theytaz et al., 2014). It has been proposed that fructose is harmful because it is converted to fructose-1-phosphate by fructokinase, accelerating glycolysis by evading the rate-limiting steps of glycolysis, and promoting lipogenesis by generating dihydroxyacetone phosphate. Although this appeared plausible when fructose was believed to be metabolized in the liver, it became less credible as fructose was shown to be cleared in the small intestine (Jang et al., 2018). Overconsumption of fructose has been shown to cause its leakage to the liver (Jang et al., 2018); however, such small amount of fructose would not fully account for its metabolic toxicity. Our results provide an alternative explanation for the toxicity of fructose. Glucose is a poor substrate for the polyol pathway as the Km value for AR is 70 to 150 mM (Gabbay, 1973). Therefore, only very small amounts of fructose would be converted from glucose through the polyol pathway under normal feeding conditions. A direct inflow of fructose to the liver could mislead the cells into responding as if there has been very high level of glucose consumption, causing them to over-activate metabolic responses to glucose uptake, including lipogenesis. This hypothesis suggests that directly ingesting fructose could be particularly deleterious to human health due to a role in triggering metabolic shifts even at low levels.

Fructose is also implicated in cancer development. Feeding mice with high-fructose corn syrup enhances tumor growth independently of obesity and metabolic syndrome (Goncalves et al., 2019). Interestingly, expression levels of the polyol pathway enzyme *AR* correlate with the epithelial-to-mesenchymal transition (EMT) status in cancer cell lines as well as in cancers in patients. Moreover, knockdown of *AR* or sorbitol dehydrogenase (*SORD)* genes can block EMT *in vitro* (Schwab et al., 2018). These observations suggest that the polyol pathway links sugar metabolism to cancer metastasis. Our work lays the foundation for further important studies uncovering the molecular mechanisms linking abnormal sugar metabolism and disease development.

## Acknowledgements

We thank Shu Kondo, and the Vienna Drosophila Resource Center, NIG Fly in the National Institute of Genetics in Japan for supplying the plasmid and fly stocks. We also acknowledge Masayo Yamane, Takumi Ichikawa, Shingo Usuki for their technical assistance, and Daria Siekhaus for critical reading of the manuscript. This work was supported by a Grant-in-Aid for Scientific Research (C) from JSPS, the Program of the Joint Usage/Research Center for Developmental Medicine, Institute of Molecular Embryology and Genetics, Kumamoto University, the Ichiro Kanehara Foundation for the Promotion of Medical Sciences and Medical Care, and MEXT-Supported Program for the Strategic Research Foundation at Private Universities.

## Author Contributions

Conceptualization, H.S.; Investigation, H.S., M.Y., H.N., and T.N.; Writing-Original Draft, H.S.; Writing-Review & Editing, H.S. and A.N.; Funding Acquisition, H.S., M.K.; Resources, H.S., A.N., and M.K.; Supervision, H.S. and A.N.

## Materials and Methods

### Fry strains and dietary conditions

The following fly stocks were used: *Oregon-R*, *w, y w*, *Gr43a^GAL4^* (Miyamoto et al., 2012), *cg-GAL4* (Asha et al., 2003; Rewitz et al., 2010), *UAS-Mondo RNAi* (#109821: VDRC), *UAS-sugarbabe RNAi* (#3850R-3: NIG). *CG6084^10-1^, CG10638^10-1^, sodh1^14-1^, sodh1^9-3^* were generated using the CRISPR/Cas9 system (see below). Flies were raised at 25°C on regular fly food containing (per liter) 40 g yeast extract, 50 g cornmeal, 30 g rice bran, 100 g glucose, and 6 g agar.

### Mutagenesis and Venus knockin

The polyol mutants were generated using the CRISPR/Cas9 system as described in (Gokcezade et al., 2014). The following sgRNA targets were used for the mutagenesis of the genes encoding AR and Sodh. Breakpoints of the mutants were determined as described previously (**Figures S1–S3**) (Kina et al., 2019; Sano et al., 2015). *CG6084* and *CG10638* are doubly mutated in *AR* mutants. *Sodh-1* and *Sodh-2* are doubly mutated in *Sodh* mutants.

*CG6084*: 5’-CCCCAAGGGTCAGGTCACCG
*CG10638*: 5’-GGCTACGAGATGCCAATTCT
S*odh-1*: 5’-GATGTACACTACCTTGCACA
S*odh-2*: 5’-GTGGGCAAGGTAGTGCACGT

The knockin of Venus at the C-terminus of the Mondo coding region was performed using the CRISPR/Cas9 system. The knockin vector was constructed by combining PCR-amplified left arm and right arm fragments for recombination, and the Esp3I fragment of the pPVxRF3 vector (a gift from S. Kondo) containing Venus and 3xP3-dsRed-Express2 using NEBuilder HiFi DNA Assembly Master Mix (NEB). The combined fragment was cloned into pBluescript. The oligonucleotides used are as follows:

L-arm forward: 5’-GCTTGATATCGAATTCTGAACGACTGGAAATTTTGG
L-arm reverse: 5’-AGTTGGGGGCGTAGGGGGTGCATGCAGATTTGG
R-arm forward: 5’-TAGTATAGGAACTTCCGTTGATGCTGATGTCCTTG
R-arm reverse: 5’-CGGGCTGCAGGAATTCGAAAATGAGAGAAGATGGCGTA

The knockin vector was injected into *y w* embryo together with the sgRNA plasmid. The following sgRNA target was used.

5’-GGCCAGCATCCAAATCTGCA

### Quantitative RT-PCR

Quantitative RT-PCR was performed as described as in (Sano et al., 2015). The following primers were used:

*CCHa2* forward: 5’-GCCTACGGTCATGTGTGCTAC
*CCHa2* reverse: 5’-ATCATGGGCAGTAGGCCATT
*rp49* forward: 5’-AGTATCTGATGCCCAACATCG
*rp49* reverse: 5’-CAATCTCCTTGCGCTTCTTG

### Metabolite extraction and quantification

Ten third-instar larvae were collected, rinsed with water, and dried on the filter paper. The cuticle was torn by forceps to release the hemolymph on a Parafilm membrane. 1 μl of hemolymph was collected and immediately quenched by mixing with 300 μl of cold methanol. The samples were further mixed with 200 μl of methanol, 200 μl of H_2_O, and 200 μl of CHCl_3_ and vortexed for 20 min at room temperature. The samples were centrifuged at 20,000 *g* for 15 min at 4°C. The supernatant was mixed with 350 μl of H_2_O and vortexed for 10 min at room temperature. The samples were centrifuged at 20,000 *g* for 15 min at 4°C. The aqueous phase was collected and dried down in a vacuum concentrator. Methoxyamine pyridine solution [20 mg/ml methoxyamine hydrochloride (Wako) in pyridine] was added to the dried residue to re-dissolve and oximated for 90 min at 30°C. Then, MSTFA + 1%TMCS (Thermo) was added for trimethylsilylation for 60 min at 37°C.

The derivatized metabolites were analyzed by an Agilent 7890B GC coupled to a 5977A Mass Selective Detector (Agilent Technologies) under the following conditions: carrier gas, helium; flow rate, 0.8 ml/min; column, DB-5MS + DG (30 m × 0.25 mm, 0.25 μm film thickness; Agilent Technologies); injection mode, 1:10 split; inlet temperature, 250°C; ion source temperature, 230°C; quadrupole temperature, 150°C. The column temperature was held at 60°C for 1 min, and then increased to 325°C at a rate of 10°C/min. The detector was operated in the electron impact ionization mode. The Agilent-Fiehn GC/MS Metabolomics RTL Library was used for metabolite identification (Kind et al., 2009). Metabolites were detected by SIM mode and the peak area of interests were analyzed by QuantAnalysis software (Agilent Technologies).

### RNA-sequencing

Early third-instar larvae (72 AEL) were starved for 18 hours on water agar plates. The larvae were then re-fed on agar plates containing 10% of indicated sugar for 6 hours. Sugar plates were supplemented with 1% Brilliant Blue to visualize sugar uptake. Total RNA from whole larvae was extracted using the PureLink RNA Mini Kit (Life Technologies). The library was constructed using the TruSeq Stranded mRNA LT Sample Prerp Kit (Illmina). RNA-seq was performed with NextSeq 500 (Illumina), targeting at least 14 million, single-end reads of 75 bp in size. The quality of the reads was assessed using FastQC (version 0.11.5). The reads were mapped to the FlyBase reference genome (Dmel Release 6.19) using Tophat2 (Kim et al., 2013). Transcript abundance and splice variant identification were determined using Cufflinks (Trapnell et al., 2010), and differential expression analysis was performed using CuffDiff (Trapnell et al., 2010).

### Histochemistry

Larval fat bodies were dissected and fixed with 4% paraformaldehyde in PBS for 30 minutes. Mondo::Venus was detected with rabbit anti-GFP antibody (Thermo Fisher Scientific, 1: 1,000) and Alexa Fluor 488-conjugated anti-rabbit-IgG (Thermo Fisher Scientific, 1: 500). Nuclei and cortical actin were labelled with DAPI (Thermo Fisher Scientific, 1 μg/mL) and Rhodamine-conjugated phalloidin (Thermo Fisher Scientific, 1: 100), respectively. After staining, fat bodies were mounted in Vectashield Mounting Medium (Vector Labs) and imaged with TCS SP8 confocal microscope using a Plan-Apochromat 63x oil-immersion objectives (Leica Microsystems) or Fluoview FV1000 confocal microscope using a UPlanSApo 60x water-immersion objectives (Olympus).

### Image analysis

Images of the fat body were analyzed using the ImageJ2 software (version 2.0.0-rc-43, NIH). Signal intensities in nuclei and whole cell were measured and the ratio of nuclear to whole cell signals was calculated on the same section.

### Statistics

The data in each graph were presented as means ± SEM. Two-tailed t-test was used to evaluate the significance of the results between two samples. For multiple comparisons, Dunnett test was used. A significance level of p<0.05 was used for all tests, which was marked by an asterisk in the figures.

## Supplemental Information

**Figure S1.**
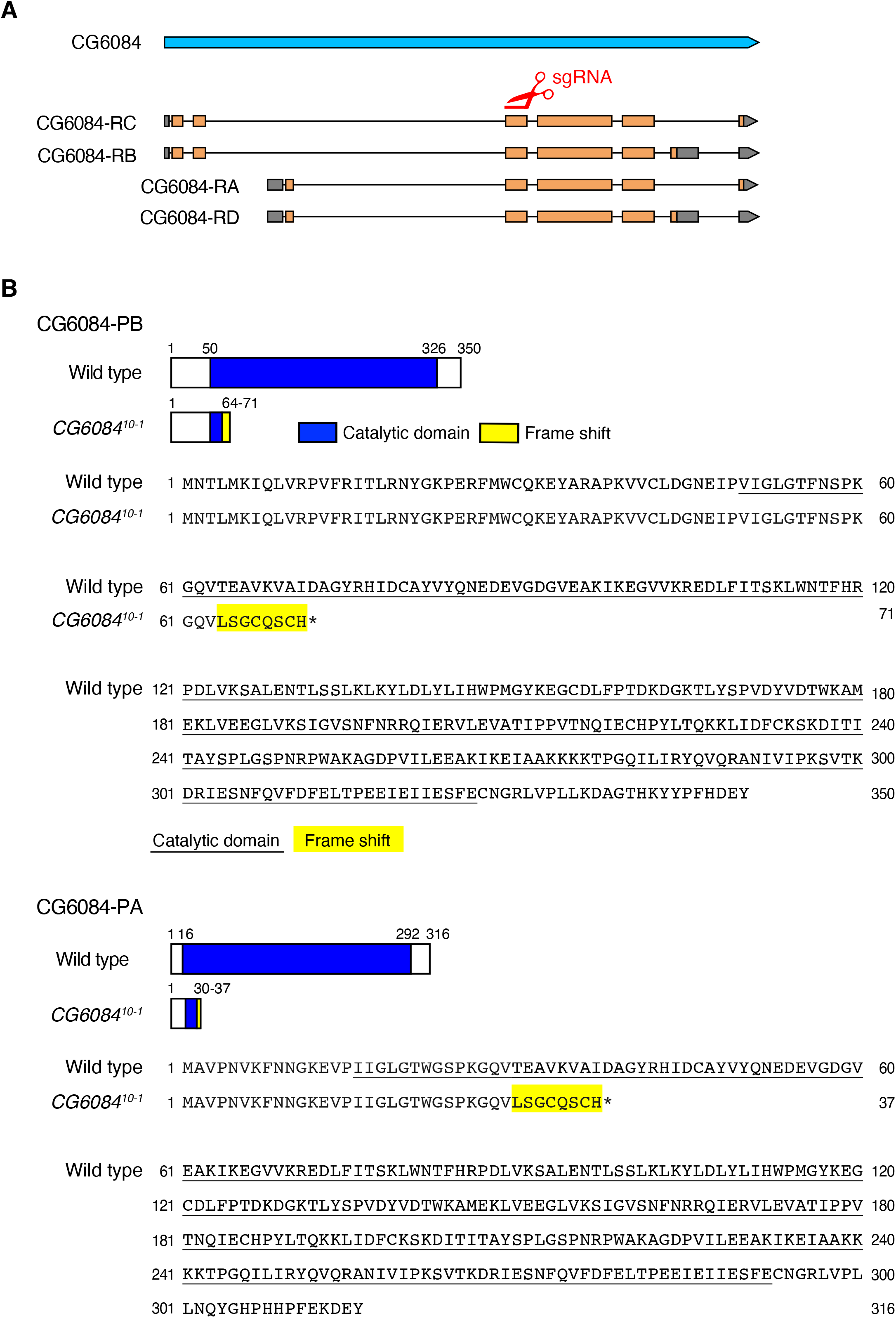
Generation of the *CG6084* mutant allele. (A) CRISPR-mediated mutagenesis of *CG6084.* A sgRNA was designed against the sequence in the exon common to all *CG6084* isoforms. (B) Breakpoint of the *CG6084^10-1^* allele. The *CG6084^10-1^* mutation caused a frame-shift (yellow) leading to a premature termination in all isoforms of the CG6084 protein. The mutant proteins lack most of the catalytic domain of the CG6084 protein (blue in schematic, underlined in sequence).

**Figure S2.**
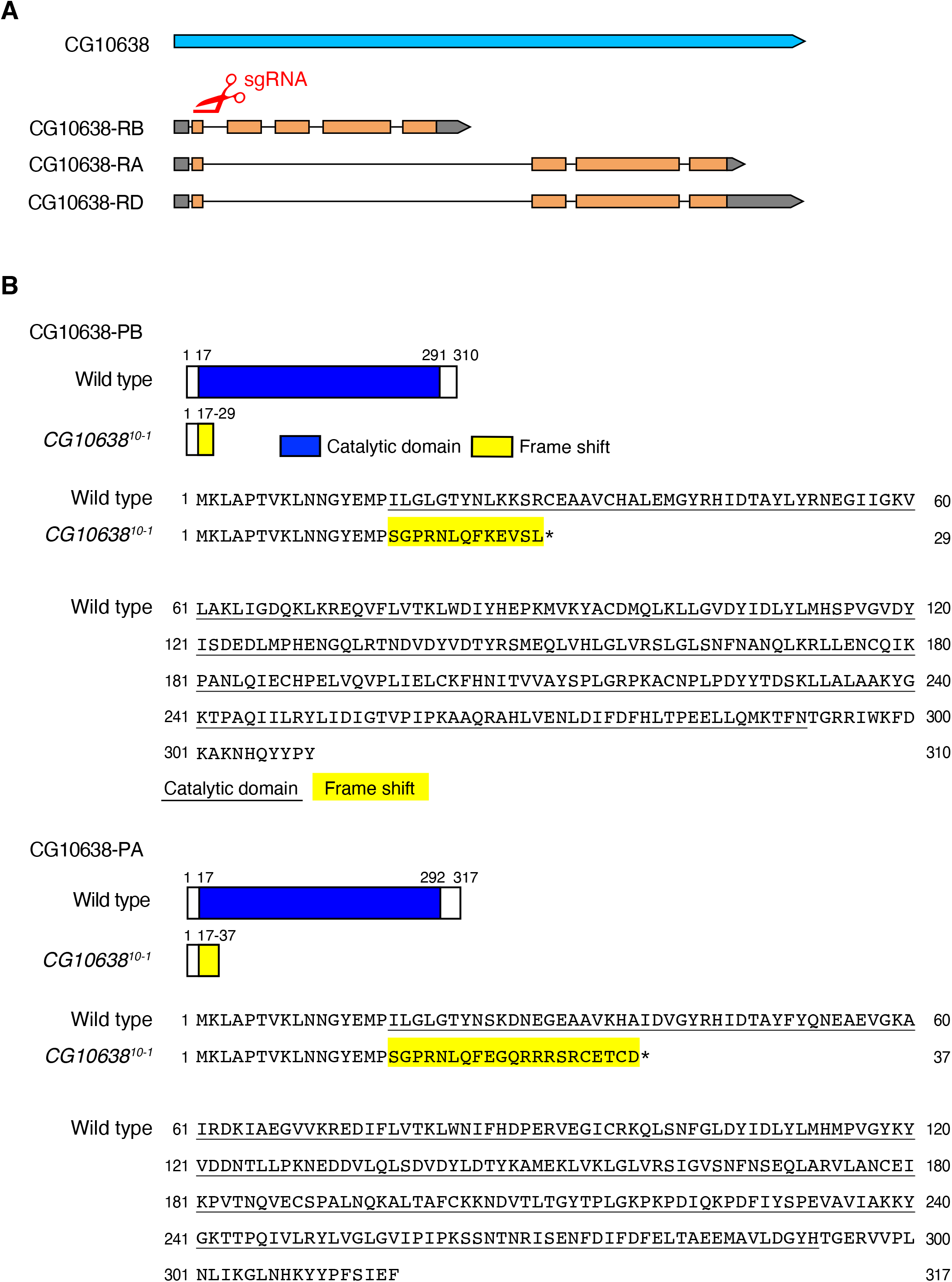
Generation of the *CG10638* mutant allele. (A) CRISPR-mediated mutagenesis of *CG10638.* A sgRNA was designed against the sequence in the common exon of *CG10638* isoforms. (B) Breakpoint of the *CG10638^10-1^* allele. The *CG10638^10-1^* mutation caused a frame-shift (yellow) leading to premature termination of all isoforms of the CG10638 protein. The mutant proteins lack most of the catalytic domain of the CG10638 protein (blue in schematic, underlined in sequence).

**Figure S3.**
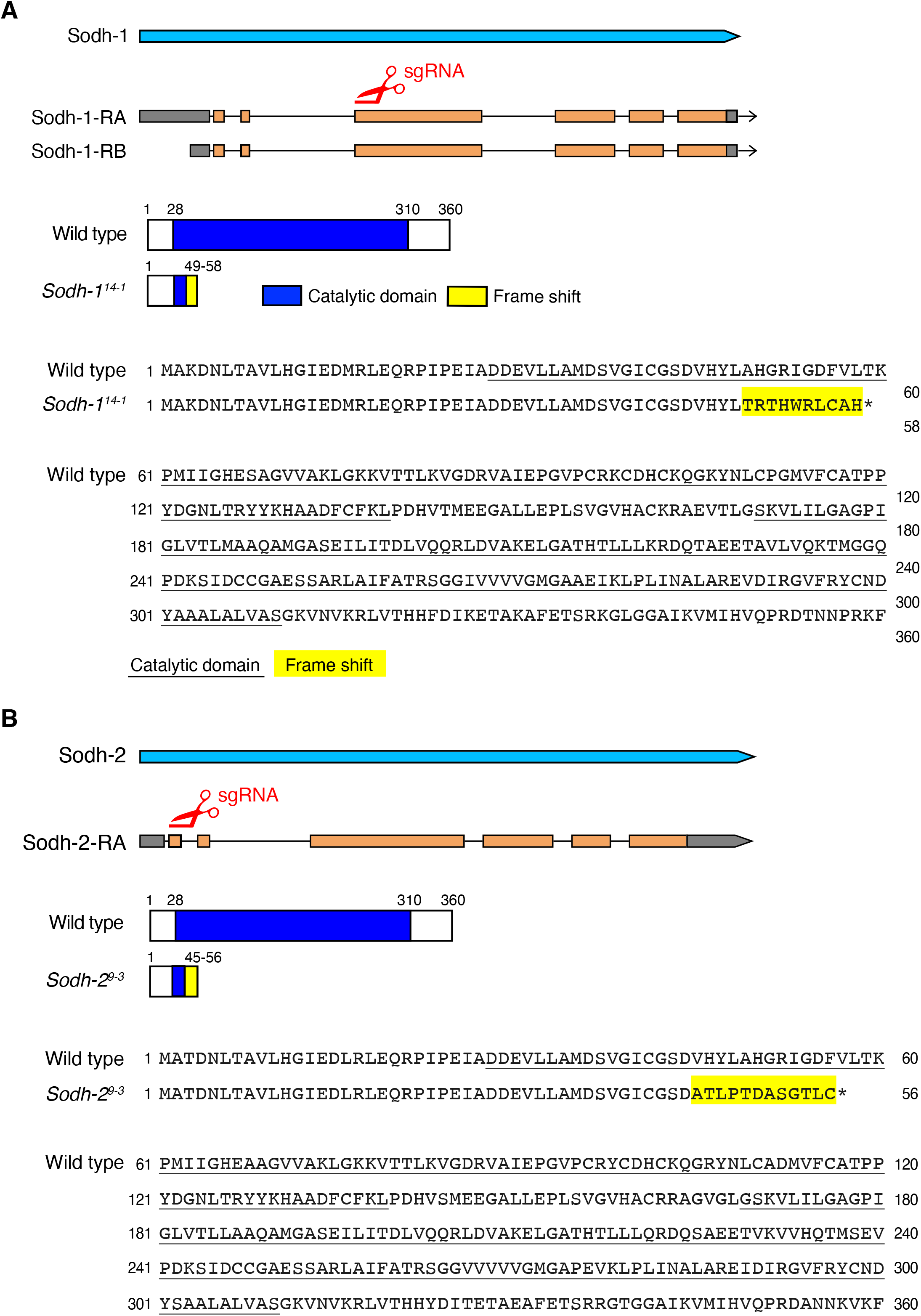
Generation of the *Sodh* mutant alleles. (A) CRISPR-mediated mutagenesis of *Sodh-1.* A sgRNA was designed against the sequence in the exon common to all *Sodh-1* isoforms. The *Sodh-1^14-1^* mutation caused a frame-shift (yellow) leading to premature termination of both isoforms of the Sodh-1 protein. The mutant proteins lack most of the catalytic domain of the Sodh-1 protein (blue in schematic, underlined in sequence). (B) CRISPR-mediated mutagenesis of *Sodh-2.* A sgRNA was designed against the sequence in the first exon of *Sodh-2.* The *Sodh-2^9-3^* mutation caused a frame-shift (yellow) leading to premature termination of the Sodh-2 protein. The mutant proteins lack most of the catalytic domain of the Sodh-2 protein (blue in schematic, underlined in sequence).

**Figure S4.**
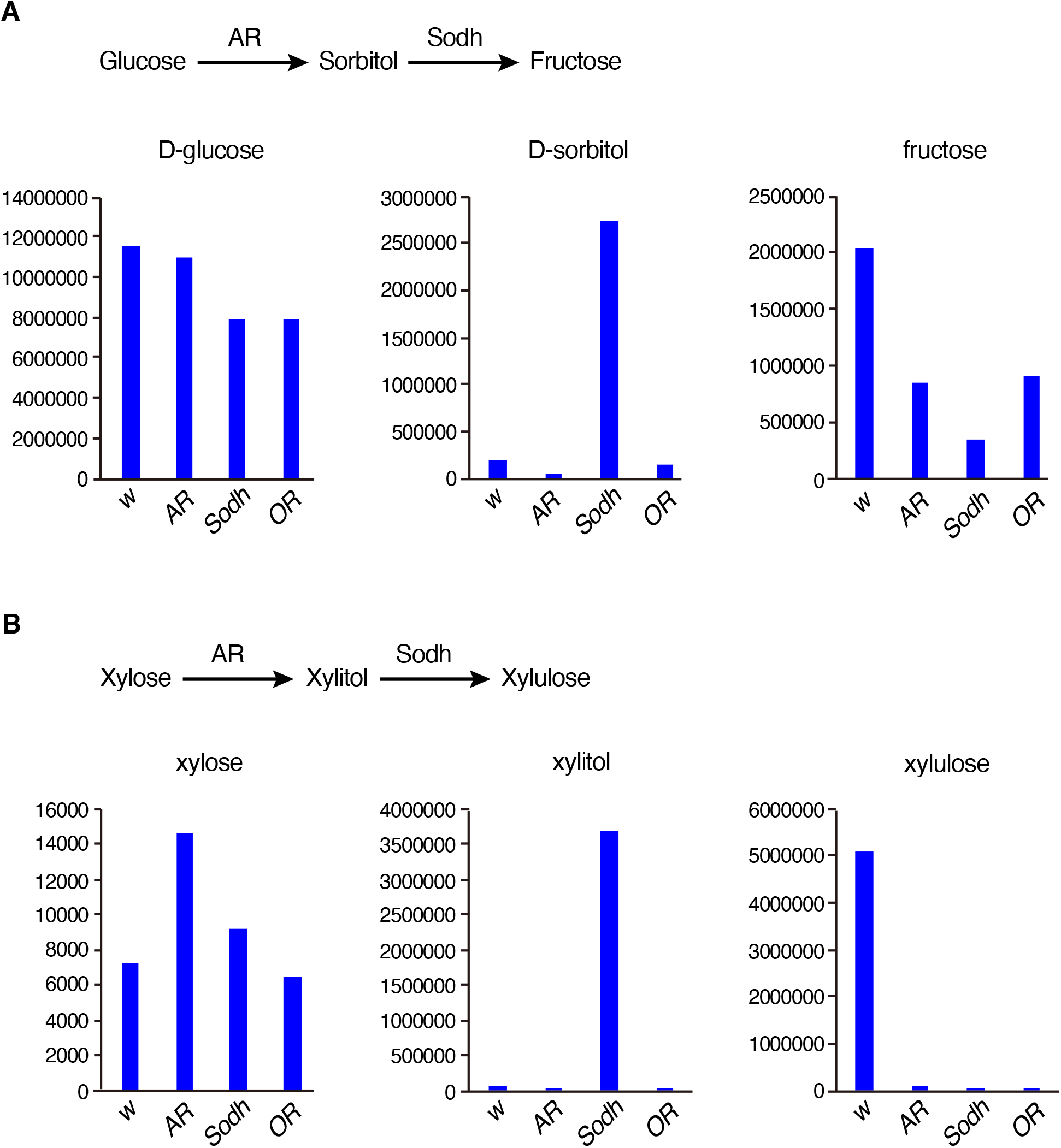
Metabolic phenotype of *AR* and *Sodh* mutants. (A) The amount of glucose, sorbitol, and fructose contained in the hemolymph of *AR* and *Sodh* mutant third-instar larvae (72 AEL) was measured using GS/MS. (B) The amount of xylose, xylitol, and xylulose contained in the hemolymph of *AR* and *Sodh* mutant third-instar larvae (72 AEL) was measured using GS/MS. *White (w)* and *Oregon-R (OR)* were used as a control.

**Figure S5.**
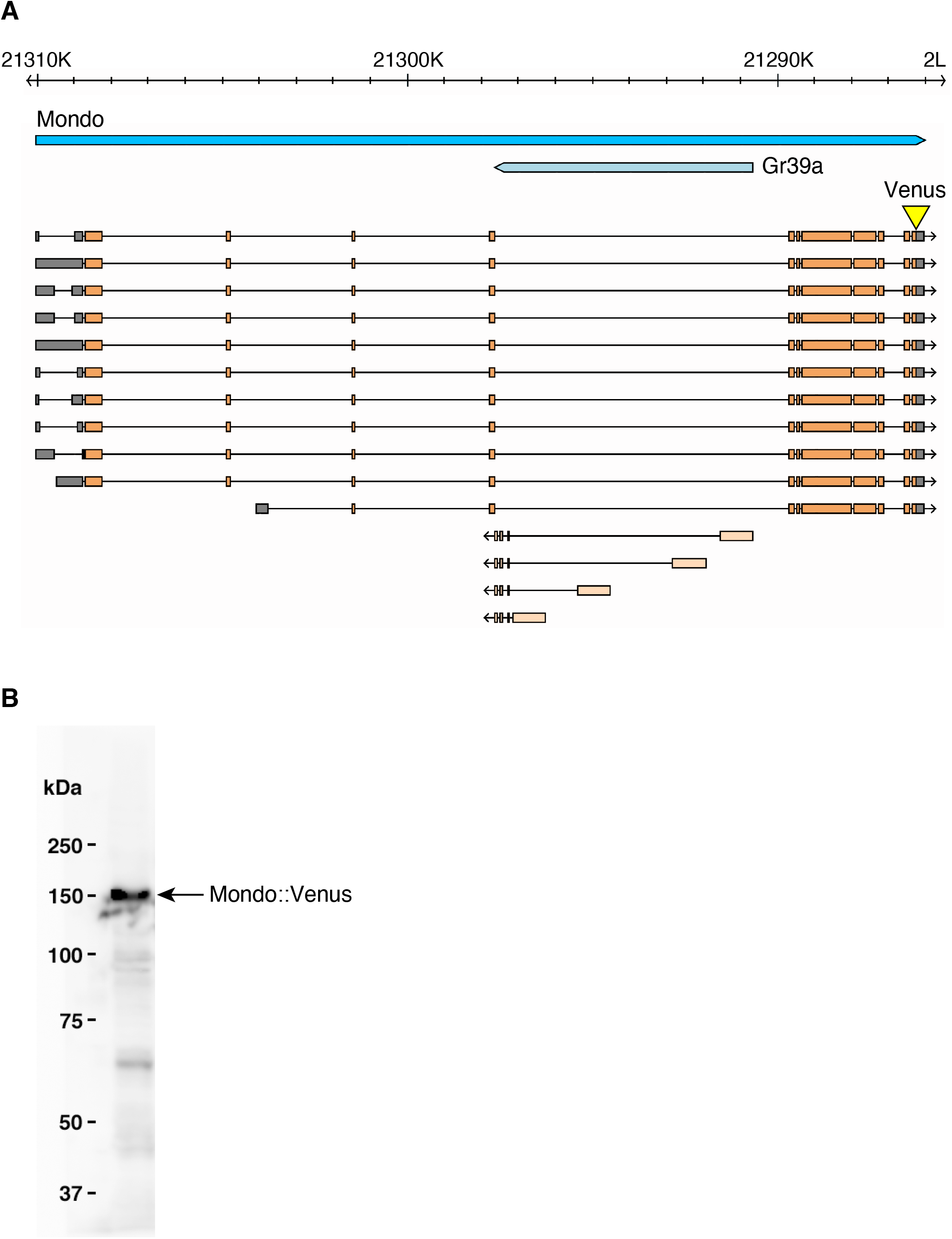
Knockin of the Venus fluorescent protein in the *Mondo* locus. (A) Schematic drawing of the Mondo locus (adapted from FlyBase, http://flybase.org). The Venus fluorescent protein was knocked-in at the C-terminus of the Mondo coding region (yellow). (B) Western blot using fat body extracts from the Mondo::Venus line. The Mondo::Venus fusion protein was detected using the anti-GFP polyclonal antibody.

**Figure S6.**
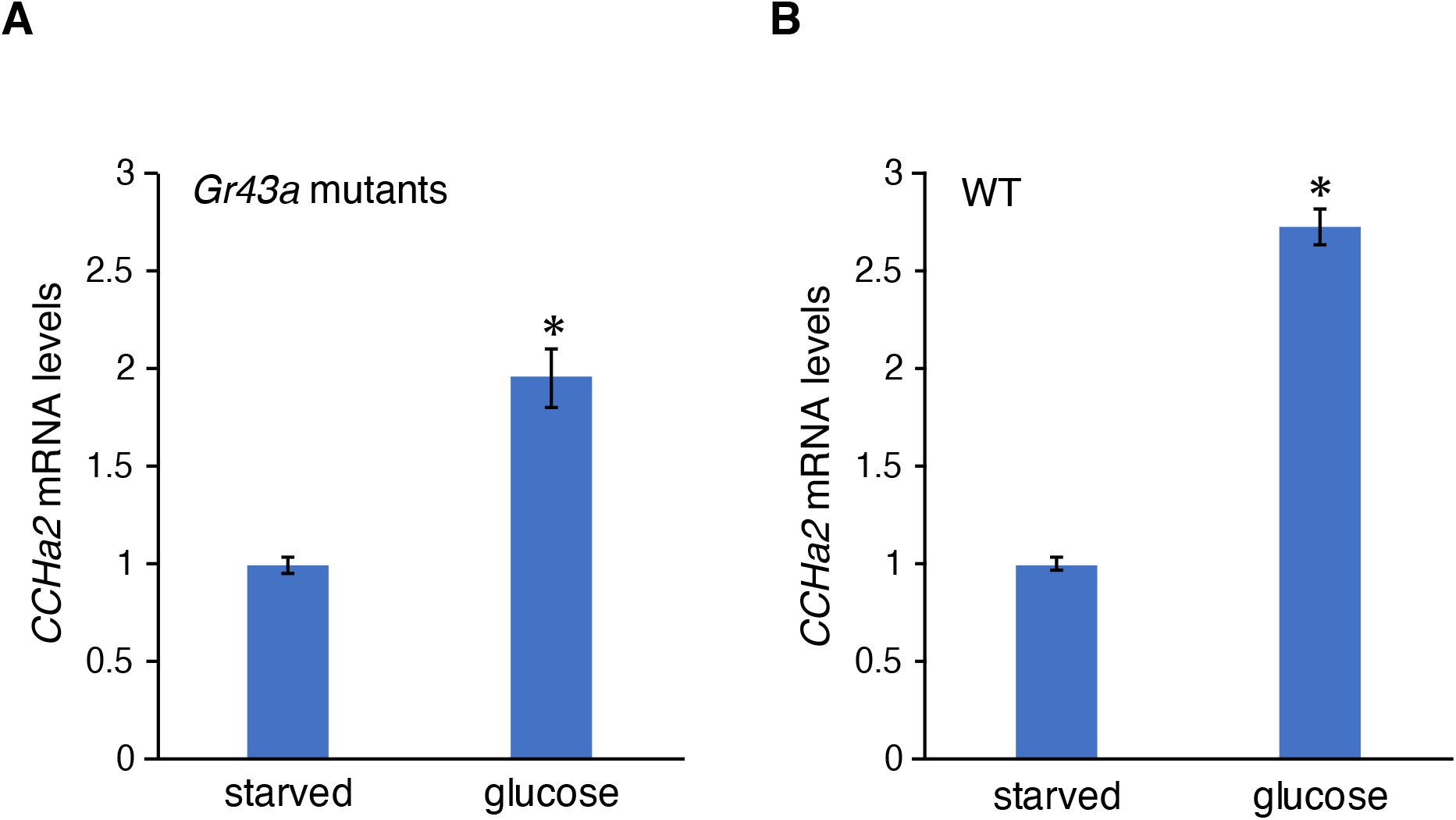
*Gr43a* is dispensable for glucose-dependent *CCHa2* expression. Glucose-dependent *CCHa2* expression was tested in *Gr43a* mutants. Third-instar larvae (72 hours AEL) starved for 18 hours were re-fed with glucose for 6 hours, and *CCHa2* mRNA levels in whole animals were measured by RT-qPCR. Induction of *CCHa2* expression in *Gr43a* mutants (A) was comparable to that in wild type (B).

## References

Agrawal, N., Delanoue, R., Mauri, A., Basco, D., Pasco, M., Thorens, B., and Leopold, P. (2016). The Drosophila TNF Eiger Is an Adipokine that Acts on Insulin-Producing Cells to Mediate Nutrient Response. Cell Metab 23, 675–684.

Arden, C., Tudhope, S.J., Petrie, J.L., Al-Oanzi, Z.H., Cullen, K.S., Lange, A.J., Towle, H.C., and Agius, L. (2012). Fructose 2,6-bisphosphate is essential for glucose-regulated gene transcription of glucose-6-phosphatase and other ChREBP target genes in hepatocytes. Biochem J 443, 111–123.

Asha, H., Nagy, I., Kovacs, G., Stetson, D., Ando, I., and Dearolf, C.R. (2003). Analysis of Ras-induced overproliferation in Drosophila hemocytes. Genetics 163, 203–215.

Bray, G.A., Nielsen, S.J., and Popkin, B.M. (2004). Consumption of high-fructose corn syrup in beverages may play a role in the epidemic of obesity. Am J Clin Nutr 79, 537–543.

Brownlee, M. (2001). Biochemistry and molecular cell biology of diabetic complications. Nature 414, 813–820.

Chintapalli, V.R.W., J.; Herzyk, P.; Dow, J.A.T. (2010). FlyAtlas: survey of adult and larval expression. Personal communication to FlyBase: http://www.flyatlasorg/.

Davies, M.N., O’Callaghan, B.L., and Towle, H.C. (2010). Activation and repression of glucose-stimulated ChREBP requires the concerted action of multiple domains within the MondoA conserved region. Am J Physiol Endocrinol Metab 299, E665–674.

Delanoue, R., Meschi, E., Agrawal, N., Mauri, A., Tsatskis, Y., McNeill, H., and Leopold, P. (2016). Drosophila insulin release is triggered by adipose Stunted ligand to brain Methuselah receptor. Science 353, 1553–1556.

Dentin, R., Tomas-Cobos, L., Foufelle, F., Leopold, J., Girard, J., Postic, C., and Ferre, P. (2012). Glucose 6-phosphate, rather than xylulose 5-phosphate, is required for the activation of ChREBP in response to glucose in the liver. J Hepatol 56, 199–209.

Diaz-Moralli, S., Ramos-Montoya, A., Marin, S., Fernandez-Alvarez, A., Casado, M., and Cascante, M. (2012). Target metabolomics revealed complementary roles of hexose- and pentose-phosphates in the regulation of carbohydrate-dependent gene expression. Am J Physiol Endocrinol Metab 303, E234–242.

Fagerberg, L., Hallstrom, B.M., Oksvold, P., Kampf, C., Djureinovic, D., Odeberg, J., Habuka, M., Tahmasebpoor, S., Danielsson, A., Edlund, K., et al. (2014). Analysis of the human tissue-specific expression by genome-wide integration of transcriptomics and antibody-based proteomics. Mol Cell Proteomics 13, 397–406.

Fridell, Y.W., Hoh, M., Kreneisz, O., Hosier, S., Chang, C., Scantling, D., Mulkey, D.K., and Helfand, S.L. (2009). Increased uncoupling protein (UCP) activity in Drosophila insulin-producing neurons attenuates insulin signaling and extends lifespan. Aging (Albany NY) 1, 699–713.

Gabbay, K.H. (1973). The sorbitol pathway and the complications of diabetes. N Engl J Med 288, 831–836.

Gokcezade, J., Sienski, G., and Duchek, P. (2014). Efficient CRISPR/Cas9 plasmids for rapid and versatile genome editing in Drosophila. G3 (Bethesda) 4, 2279–2282.

Goncalves, M.D., Lu, C., Tutnauer, J., Hartman, T.E., Hwang, S.K., Murphy, C.J., Pauli, C., Morris, R., Taylor, S., Bosch, K., et al. (2019). High-fructose corn syrup enhances intestinal tumor growth in mice. Science 363, 1345–1349.

Graveley, B.R.M.G.,; Brooks, A.N.; Carlson, J.W.; Cherbas, L.; Davis, C.A.; Duff, M.; Eads, B.; Landolin, J.; Sandler, J.; Wan, K.H.; Andrews, J.; Brenner, S.E.; Cherbas, P.; Gingeras, T.R.; Hoskins, R.; Kaufman, T.; Celniker, S.E. (2011). The D. melanogaster transcriptome: modENCODE RNA-Seq data for dissected tissues: http://www.modencode.org/celniker/.

Havula, E., Teesalu, M., Hyotylainen, T., Seppala, H., Hasygar, K., Auvinen, P., Oresic, M., Sandmann, T., and Hietakangas, V. (2013). Mondo/ChREBP-Mlx-regulated transcriptional network is essential for dietary sugar tolerance in Drosophila. PLoS Genet 9, e1003438.

Hers, H.G. (1956). [The mechanism of the transformation of glucose in fructose in the seminal vesicles]. Biochim Biophys Acta 22, 202–203.

Iizuka, K., Wu, W., Horikawa, Y., and Takeda, J. (2013). Role of glucose-6-phosphate and xylulose-5-phosphate in the regulation of glucose-stimulated gene expression in the pancreatic beta cell line, INS-1E. Endocr J 60, 473–482.

Jang, C., Hui, S., Lu, W., Cowan, A.J., Morscher, R.J., Lee, G., Liu, W., Tesz, G.J., Birnbaum, M.J., and Rabinowitz, J.D. (2018). The Small Intestine Converts Dietary Fructose into Glucose and Organic Acids. Cell Metab 27, 351–361 e353.

Kabashima, T., Kawaguchi, T., Wadzinski, B.E., and Uyeda, K. (2003). Xylulose 5-phosphate mediates glucose-induced lipogenesis by xylulose 5-phosphate-activated protein phosphatase in rat liver. Proc Natl Acad Sci U S A 100, 5107–5112.

Kanehisa, M., and Goto, S. (2000). KEGG: kyoto encyclopedia of genes and genomes. Nucleic Acids Res 28, 27–30.

Kanehisa, M., Sato, Y., Furumichi, M., Morishima, K., and Tanabe, M. (2019). New approach for understanding genome variations in KEGG. Nucleic Acids Res 47, D590–D595.

Kawaguchi, T., Takenoshita, M., Kabashima, T., and Uyeda, K. (2001). Glucose and cAMP regulate the L-type pyruvate kinase gene by phosphorylation/dephosphorylation of the carbohydrate response element binding protein. Proc Natl Acad Sci U S A 98, 13710–13715.

Kim, D., Pertea, G., Trapnell, C., Pimentel, H., Kelley, R., and Salzberg, S.L. (2013). TopHat2: accurate alignment of transcriptomes in the presence of insertions, deletions and gene fusions. Genome Biol 14, R36.

Kim, J., and Neufeld, T.P. (2015). Dietary sugar promotes systemic TOR activation in Drosophila through AKH-dependent selective secretion of Dilp3. Nat Commun 6, 6846.

Kina, H., Yoshitani, T., Hanyu-Nakamura, K., and Nakamura, A. (2019). Rapid and efficient generation of GFP-knocked-in Drosophila by the CRISPR-Cas9-mediated genome editing. Dev Growth Differ 61, 265–275.

Kind, T., Wohlgemuth, G., Lee, D.Y., Lu, Y., Palazoglu, M., Shahbaz, S., and Fiehn, O. (2009). FiehnLib: mass spectral and retention index libraries for metabolomics based on quadrupole and time-of-flight gas chromatography/mass spectrometry. Anal Chem 81, 10038–10048.

Koyama, T., and Mirth, C.K. (2016). Growth-Blocking Peptides As Nutrition-Sensitive Signals for Insulin Secretion and Body Size Regulation. PLoS Biol 14, e1002392.

Li, M.V., Chang, B., Imamura, M., Poungvarin, N., and Chan, L. (2006). Glucose-dependent transcriptional regulation by an evolutionarily conserved glucose-sensing module. Diabetes 55, 1179–1189.

Li, M.V., Chen, W., Harmancey, R.N., Nuotio-Antar, A.M., Imamura, M., Saha, P., Taegtmeyer, H., and Chan, L. (2010). Glucose-6-phosphate mediates activation of the carbohydrate responsive binding protein (ChREBP). Biochem Biophys Res Commun 395, 395–400.

Lim, J.S., Mietus-Snyder, M., Valente, A., Schwarz, J.M., and Lustig, R.H. (2010). The role of fructose in the pathogenesis of NAFLD and the metabolic syndrome. Nat Rev Gastroenterol Hepatol 7, 251–264.

Lorenzi, M. (2007). The polyol pathway as a mechanism for diabetic retinopathy: attractive, elusive, and resilient. Exp Diabetes Res 2007, 61038.

Marriott, B.P., Cole, N., and Lee, E. (2009). National estimates of dietary fructose intake increased from 1977 to 2004 in the United States. J Nutr 139, 1228S–1235S.

Matsuda, H., Yamada, T., Yoshida, M., and Nishimura, T. (2015). Flies without trehalose. J Biol Chem 290, 1244–1255.

Mattila, J., Havula, E., Suominen, E., Teesalu, M., Surakka, I., Hynynen, R., Kilpinen, H., Vaananen, J., Hovatta, I., Kakela, R., et al. (2015). Mondo-Mlx Mediates Organismal Sugar Sensing through the Gli-Similar Transcription Factor Sugarbabe. Cell Rep 13, 350–364.

Mishra, D., Miyamoto, T., Rezenom, Y.H., Broussard, A., Yavuz, A., Slone, J., Russell, D.H., and Amrein, H. (2013). The molecular basis of sugar sensing in Drosophila larvae. Curr Biol 23, 1466–1471.

Miyamoto, T., Slone, J., Song, X., and Amrein, H. (2012). A fructose receptor functions as a nutrient sensor in the Drosophila brain. Cell 151, 1113–1125.

Mor, I., Cheung, E.C., and Vousden, K.H. (2011). Control of glycolysis through regulation of PFK1: old friends and recent additions. Cold Spring Harb Symp Quant Biol 76, 211–216.

Pasco, M.Y., and Leopold, P. (2012). High sugar-induced insulin resistance in Drosophila relies on the lipocalin Neural Lazarillo. PLoS One 7, e36583.

Peeters, K., Van Leemputte, F., Fischer, B., Bonini, B.M., Quezada, H., Tsytlonok, M., Haesen, D., Vanthienen, W., Bernardes, N., Gonzalez-Blas, C.B., et al. (2017). Fructose-1,6-bisphosphate couples glycolytic flux to activation of Ras. Nat Commun 8, 922.

Peterson, C.W., Stoltzman, C.A., Sighinolfi, M.P., Han, K.S., and Ayer, D.E. (2010). Glucose controls nuclear accumulation, promoter binding, and transcriptional activity of the MondoA-Mlx heterodimer. Mol Cell Biol 30, 2887–2895.

Petrie, J.L., Al-Oanzi, Z.H., Arden, C., Tudhope, S.J., Mann, J., Kieswich, J., Yaqoob, M.M., Towle, H.C., and Agius, L. (2013). Glucose induces protein targeting to glycogen in hepatocytes by fructose 2,6-bisphosphate-mediated recruitment of MondoA to the promoter. Mol Cell Biol 33, 725–738.

Rajan, A., and Perrimon, N. (2012). Drosophila cytokine unpaired 2 regulates physiological homeostasis by remotely controlling insulin secretion. Cell 151, 123–137.

Rewitz, K.F., Yamanaka, N., and O’Connor, M.B. (2010). Steroid hormone inactivation is required during the juvenile-adult transition in Drosophila. Dev Cell 19, 895–902.

Richards, P., Ourabah, S., Montagne, J., Burnol, A.F., Postic, C., and Guilmeau, S. (2017). MondoA/ChREBP: The usual suspects of transcriptional glucose sensing; Implication in pathophysiology. Metabolism 70, 133–151.

Sakiyama, H., Wynn, R.M., Lee, W.R., Fukasawa, M., Mizuguchi, H., Gardner, K.H., Repa, J.J., and Uyeda, K. (2008). Regulation of nuclear import/export of carbohydrate response element-binding protein (ChREBP): interaction of an alpha-helix of ChREBP with the 14-3-3 proteins and regulation by phosphorylation. J Biol Chem 283, 24899–24908.

Sano, H. (2015). Coupling of growth to nutritional status: The role of novel periphery-to-brain signaling by the CCHa2 peptide in Drosophila melanogaster. Fly (Austin) 9, 183–187.

Sano, H., Nakamura, A., Texada, M.J., Truman, J.W., Ishimoto, H., Kamikouchi, A., Nibu, Y., Kume, K., Ida, T., and Kojima, M. (2015). The Nutrient-Responsive Hormone CCHamide-2 Controls Growth by Regulating Insulin-like Peptides in the Brain of Drosophila melanogaster. PLoS Genet 11, e1005209.

Schwab, A., Siddiqui, A., Vazakidou, M.E., Napoli, F., Bottcher, M., Menchicchi, B., Raza, U., Saatci, O., Krebs, A.M., Ferrazzi, F., et al. (2018). Polyol Pathway Links Glucose Metabolism to the Aggressiveness of Cancer Cells. Cancer Res 78, 1604–1618.

Stanhope, K.L., Schwarz, J.M., Keim, N.L., Griffen, S.C., Bremer, A.A., Graham, J.L., Hatcher, B., Cox, C.L., Dyachenko, A., Zhang, W., et al. (2009). Consuming fructose-sweetened, not glucose-sweetened, beverages increases visceral adiposity and lipids and decreases insulin sensitivity in overweight/obese humans. J Clin Invest 119, 1322–1334.

Stoltzman, C.A., Peterson, C.W., Breen, K.T., Muoio, D.M., Billin, A.N., and Ayer, D.E. (2008). Glucose sensing by MondoA:Mlx complexes: a role for hexokinases and direct regulation of thioredoxin-interacting protein expression. Proc Natl Acad Sci U S A 105, 6912–6917.

Straub, S.G., and Sharp, G.W. (2002). Glucose-stimulated signaling pathways in biphasic insulin secretion. Diabetes Metab Res Rev 18, 451–463.

Sun, J., Liu, C., Bai, X., Li, X., Li, J., Zhang, Z., Zhang, Y., Guo, J., and Li, Y. (2017). Drosophila FIT is a protein-specific satiety hormone essential for feeding control. Nat Commun 8, 14161.

Theytaz, F., de Giorgi, S., Hodson, L., Stefanoni, N., Rey, V., Schneiter, P., Giusti, V., and Tappy, L. (2014). Metabolic fate of fructose ingested with and without glucose in a mixed meal. Nutrients 6, 2632–2649.

Trapnell, C., Williams, B.A., Pertea, G., Mortazavi, A., Kwan, G., van Baren, M.J., Salzberg, S.L., Wold, B.J., and Pachter, L. (2010). Transcript assembly and quantification by RNA-Seq reveals unannotated transcripts and isoform switching during cell differentiation. Nat Biotechnol 28, 511–515.

Tsatsos, N.G., Davies, M.N., O’Callaghan, B.L., and Towle, H.C. (2008). Identification and function of phosphorylation in the glucose-regulated transcription factor ChREBP. Biochem J 411, 261–270.

Ugrankar, R., Berglund, E., Akdemir, F., Tran, C., Kim, M.S., Noh, J., Schneider, R., Ebert, B., and Graff, J.M. (2015). Drosophila glucome screening identifies Ck1alpha as a regulator of mammalian glucose metabolism. Nat Commun 6, 7102.

